# *Drosophila* Trus, the orthologue of mammalian PDCD2L, is required for proper cell proliferation, larval developmental timing, and oogenesis

**DOI:** 10.1101/2024.10.24.620039

**Authors:** Saeko Takada, Bonnie J. Bolkan, MaryJane O’Connor, Michael Goldberg, Michael B. O’Connor

## Abstract

Toys are us (Trus) is the *Drosophila melanogaster* ortholog of mammalian Programmed Cell Death 2-Like (PDCD2L), a protein that has been implicated in ribosome biogenesis, cell cycle regulation, and oncogenesis. In this study, we examined the function of Trus during *Drosophila* development. CRISPR/Cas9 generated null mutations in *trus* lead to partial embryonic lethality, significant larval developmental delay, and complete pre-pupal lethality. In mutant larvae, we found decreased cell proliferation and growth defects in the brain and imaginal discs. Mapping relevant tissues for Trus function using *trus RNAi* and *trus* mutant rescue experiments revealed that imaginal disc defects are primarily responsible for the developmental delay, while the pre-pupal lethality is likely associated with faulty central nervous system (CNS) development. Examination of the molecular mechanism behind the developmental delay phenotype revealed that *trus* mutations induce the Xrp1-Dilp8 ribosomal stress-response in growth-impaired imaginal discs, and this signaling pathway attenuates production of the hormone ecdysone in the prothoracic gland. Additional Tap-tagging and mass spectrometry of components in Trus complexes isolated from *Drosophila* Kc cells identified Ribosomal protein subunit 2 (RpS2), which is coded by *string of pearls (sop)* in *Drosophila,* and Eukaryotic translation elongation factor 1 alpha 1 (eEF1α1) as interacting factors. We discuss the implication of these findings with respect to the similarity and differences in *trus* genetic null mutant phenotypes compared to the haplo-insufficiency phenotypes produced by heterozygosity for mutants in Minute genes and other genes involved in ribosome biogenesis.

**Authors Summary:** Ribosomes are essential macromolecular machines required for decoding mRNA to make proteins, the major biomolecules that carry out all central cellular functions. As such, their structural and operational integrity is critical to organismal survival, and mutations that disrupt proper stoichiometry or assembly of ribosomes produce serious pathological consequences during an organism’s development and/or adult life. The ribosome assembly factor PDCD2L is highly conserved from yeast to man, yet its overall function and requirement during development is poorly understood. By examining the developmental consequences of null mutations in *trus*, which encodes the *Drosophila* PDCD2L homolog, we demonstrate an essential role for this factor in cell-cycle regulation. Furthermore, disruption of Trus function in mitotically dividing imaginal tissue activates the Xrp1-dilp8 stress response pathway which limits production of ecdysone, the major arthropod molting hormone, leading to severe developmental delay during larval stages. These studies provide new insights on the requirements of this highly conserved ribosome assemble factor during development.

## Introduction

Ribosomes are fundamental macromolecular machines present in all life forms that are required for decoding the genome to build cells that have specific identities and functions. As such, their assembly is subject to strict quality control [1] and when aberrations occur, cellular dysfunction and organismal disease are a frequent outcome. In humans, defects in ribosome assembly and subunit production are collectively known as ribosomopathies and produce a myriad of pathologies including microcephaly, intellectual disability, neurodegeneration, seizures, various types of cancers and numerous additional syndromes [2].

Over one hundred years ago, *Drosophila* offered the first insight into the importance of proper ribosome biogenesis, through the isolation of haplo-insufficient *‘Minute’* mutants which, as heterozygotes (*M/+*), develop with a distinctive thin and small bristle phenotype [3]. These heterozygous mutants also exhibit developmental delay, occasional notched eyes, and reduced viability and fertility, while homozygous mutants die at early developmental stages. *M/+* mutations were subsequently shown to almost exclusively affect ribosomal protein subunits (RPS) [4–6]. Subsequent work uncovered the interesting phenomenon of cell competition whereby slow growing *M/+* cells (losers), when they are induced as “mosaic clones” in a field of wild-type cells (winners), undergo apoptosis and are eliminated [7]. Cell competition is not just confined to *M/+* mutations but is observed in similar context-dependent elimination of viable cells in *Drosophila* imaginal epithelia when there exists a discrepancy in genotype between neighboring cells such as heterozygosity for apicobasal polarity genes (*scrib, dlg*), endocytosis components (*Vps25*, *Rab5*), ER stress, and others (reviewed in Nagata and Igaki, 2024)[8]. Cell competition has also been documented in mice, zebrafish, and mammalian tissue culture cells [9–12] suggesting that it may be a universal mechanism for detecting and eliminating growth-compromised clones of cells in an otherwise healthy tissue.

While the molecular mechanism(s) responsible for the various types of cell competition are still not completely understood, in the case of *M/+* mutants, several recent studies have shown that they activate a novel stress response pathway involving RpS12-mediated induction of the transcription factor Xrp1 [13–16]. Xrp1, likely in complex with another basic leucine-zipper protein (bZIP) Irbp18 [17], then activates downstream targets including JNK-mediate apoptotic genes, DNA damage repair pathways, and antioxidant genes [18]. In addition, Xrp1 stimulates expression of protein kinase R (PKR)-like endoplasmic reticulum kinase (PERK) which then phosphorylates eukaryotic translation initiation factor 2A (EIF2A) leading to a reduction in protein translation in the loser cells and their eventual loss by apoptosis [19–21].

One of the most highly induced Xrp1-dependent genes in *M*/+ cells is *dilp8* [22]. This secreted insulin/relaxin-related factor is released from *M/+* imaginal disc cells and inhibits production of neuronally-derived PTTH, the principal neuropeptide that sets the pace of larval developmental maturation through stimulation of ecdysone production in the prothoracic gland (PG) [14–16]. Knockdown of either *Xrp1* or *dilp8* in *M/+* cells is sufficient to restore developmental timing to a near normal pace suggesting that Dilp8 activity is the primary mechanism responsible for producing delayed development of *M/+* larvae. In addition to the phenotypes caused by mutations in structural subunits of ribosomes, related phenotypes are often produced by mutations in other aspects of ribosome biogenesis.

For example, in *Drosophila, RNAi*-mediated knockdown or genetic mutations in components of the nucleolus including Nop60b, Nop140 and Noc1, or Rpl-135, a subunit of the Pol I RNA transcription complex, and Paip1, a poly A binding protein that stimulations translation initiation, can also result in reduced growth and developmental delay [23–26]. In the case of Noc1, *RNAi* mediated knockdown in the wing imaginal disc also resulted in upregulation of *dilp8* via *Xrp1* activation similar to what is seen in *Minute* mutations and likely accounts for the slow development phenotype [27].

Several other well-studied regulators of ribosome biogenesis are vertebrate 40S ribosomal protein uS5/RPS2 and its interaction partners PDCD2 and PDCD2L [28]. In *Drosophila*, the uS5/RPS2 ortholog is encoded by the *string of pearls* (*sop)/RpS2* gene, while the orthologs of PDCD2 and PDCD2L are encoded by *Zfrp8* (*<u>Z</u>inc finger protein <u>RP</u>-<u>8</u>*) and *trus,* respectively [29–31]. The moniker *sop* refers to the oogenesis defect seen in females from recessive hypomorphic sterile alleles which block oocyte development leading to a logjam accumulation of pre-oocytes within each ovariole. These mutants also exhibit classic *Minute-like* phenotypes such as thin bristles and developmental delay [29].

Loss of *uS5/RPS2* in yeast and human cell lines leads to reduction in processing of pre-20S and 21S rRNA precursors, respectively, and defects in nuclear export of pre-40S complexes [32, 33]. Biochemical pull-down experiments from yeast and human cell lines identified uS5 as a binding partner of both PDCD2 and PDCD2L [34–37]. These proteins are paralogs that appear to have arisen through gene duplication before the split of animals from plants and fungi [38]. In mice, loss of PDCD2 leads to a failure of the Inner cell mass development after implantation likely due to reduced viability and proliferation of embryonic stem cells [39], while mutants of PDCD2L develop further but die and are resorbed at around day E12.5 [40]. Studies in human cell lines and yeast suggest that PDCD2 and PDCD2L bind to common or overlapping sites on uS5 and act as either a chaperone or an adaptor to facilitate several distinct steps of pre-40S ribosomal particle assembly and transport, and thus are essential for 40S ribosomal subunit biogenesis [28, 34, 36].

In *Drosophila*, loss of Zfrp8, the PDCD2 homolog, leads to several developmental defects including delayed larval growth, lymph gland over-proliferation, and reduced germ cell proliferation followed by oocyte arrest and degeneration [30, 41, 42]. Mass spectrometry analysis of complexes containing Tap tagged Zfrp8 identified 30 potential binding partners including Nop60B and uS5/Sop, 5 other ribosomal subunits, several translational elongation and initiation factors and FMRP the fragile-X mental retardation protein [43]. The variety of complexes formed by Zfrp8 suggest that it is likely involved in numerous other molecular processes aside from acting as a chaperone of uS5. Consistent with this view, Zfrp8 is required for proper localization of FMRP in oocytes where the complex likely targets select mRNAs for repression. Interestingly, Zfrp8 and FMRP also appear to regulate heterochromatin formation and transposon de-repression; however, the molecular details for how the various Zfrp8 complexes affect this are still unclear [43].

In this report, we analyze in detail the phenotypes associated with Toys are us (Trus), the *Drosophila* ortholog of mammalian PDCD2L. We compare the various *trus* mutant phenotypes with those produced by *Minute*, *Zfrp8,* and knockdown of other ribosome assembly factors and find both striking similarities but also notable differences, suggesting that Trus loss triggers common ribosomal/proteostasis stress pathways such as Xrp1-Dilp8, but also produces distinctive phenotypes indicative of its specific role in ribosome assembly or its requirement in other biological/developmental processes.

## Results

### Production of *trus* CRISPR/Cas9 mutants

The original *trus^1^* mutant (*<u>t</u>oys a<u>r</u>e <u>us</u>*) was isolated from the Zucker EMS mutant collection [31, 44]. The mutation caused significant developmental delay throughout development and 3^rd^ instar larvae wandered up to 10 days before pupariating and subsequently dying as pre-pupae [31]. Sequencing analysis revealed that the *trus^1^* allele carries a point mutation in the start codon, which may result in a failure of translational initiation (Fig. 1A) [31]. We found that a small percentage of heterozygous *trus^1^/Dftrus* (*Df(3R)BSC847*) developed to pharates and a few enclosed as adults that showed notched eyes and thin/short bristles (Fig. S1). These phenotypes, in addition to the prolonged developmental time, resemble the haplo-insufficiency *‘Minute’* syndrome that is often observed in flies carrying a mutation in one of the genes encoding ribosomal proteins [6]. Since no adult escapers were obtained from *trus^1^* homozygous, we inferred that *trus^1^* might be an antimorph allele. This could arise from an aberrant translational start at one of the methionine codons downstream of the normal initiation codon which would produce an N-terminally truncated protein. There are four additional candidate initiation sequences which weakly align to the Kozak consensus [45, 46]. Candidate 1 is from −7 bp upstream, candidate 2 is from +381 bp downstream, candidate 3 is from +415 bp downstream, and candidate 4 is from +579 bp downstream of the original initiation start codon. Among them, candidate 2 and candidate 4 are in-frame with the Trus protein sequence and could produce N-terminally truncated 238 a.a. (∼27kDa) or 173 a.a. (∼19kDa) Trus fragments, respectively.

**Fig. 1.**
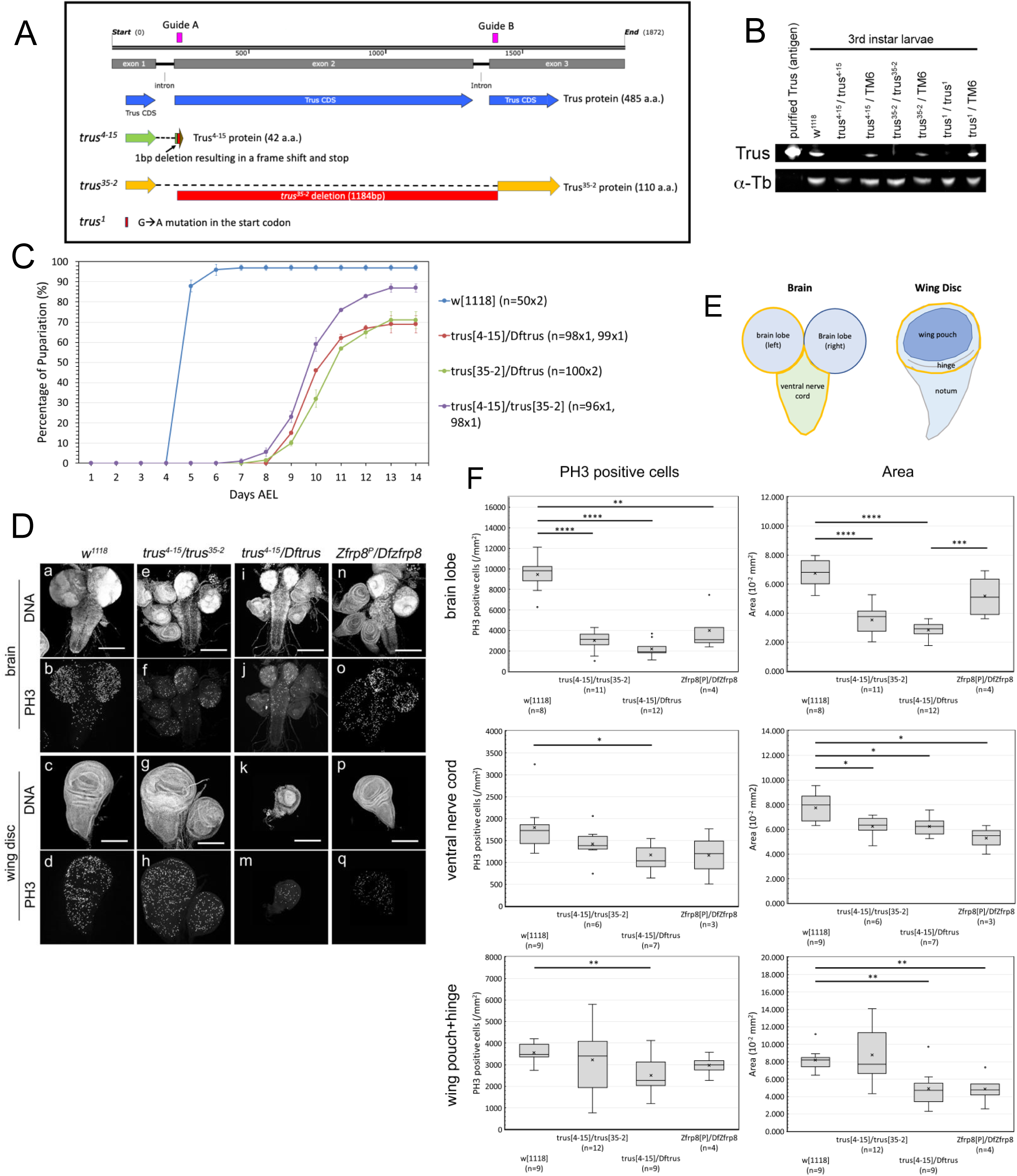
CRISPR/Cas9 induced *trus* mutations cause developmental delay and defects in tissue growth and cell proliferation during the larval stage. (A) A diagram showing *trus* genomic region and *trus* mutant alleles that are used in this study. (Blue) Trus CDS, (Magenta) two guide RNAs designed for CRISPR/Cas9 mutagenesis, (Red) *trus* mutations and deletion. (Green) *trus^4-15^* allele coding 42 a.a. fragment. (Orange) *trus^35-2^* allele including 1184 bp in-frame deletion and coding 110 a.a. Trus fragment. **(B)** Western-blot analysis of 3rd instar larval lysate of *trus* homozygotes and heterozygotes over a balancer chromosome (*TM6B P[Dfd-GMR-nvYPF] Sb*). Purified full length recombinant Trus protein used as antigen for the anti-Trus antibody production is shown on the far left. (Trus) affinity-purified anti-Trus antibody produced in this study. (α-Tb) mouse anti-α−Tubulin monoclonal antibody (DM1A) (Sigma-Aldrich T9026). **(C)** Pupariation timing of *trus* mutant and *w^1118^*. Each data point on the graph indicates an average pupariation percentage of two separate plates. The vertical line on each data point indicates the standard deviation. Numbers of 1st instar larvae picked at 1 Day AEL are shown in parentheses after the genotype. Percentage of pupariation at 14 days AEL were 97%, 87%, 69%, 71% for *w^1118^*, *trus^4-15^/trus^35-2^*, *trus^4-15^/Dftrus*, and *trus^35-2^/Dftrus*, respectively. After pupariation, none of the *trus* mutant larvae developed further and eventually died (pre-pupal lethal). More than 95% of the *w^1118^* animals eclosed as adults. **(D)** Representative images of brains and wing discs that were dissected from 3^rd^ instar wandering larvae. Genotypes indicated at the top of the panels. DNA was stained with DAPI and mitotic cells are detected with anti-phospho-HistoneH3 (PH3) antibody. Scale bar: 200μm. **(E)** Diagrams showing the larval brain and wing-disc. The areas surrounded by yellow lines indicates brain lobe, ventral nerve cord, and wing pouch plus hinge areas that are quantified in F. **(F)** Box-and-whisker plots of PH3 positive cells/mm^2^ (left column) and area in mm^2^ (right column) for brain lobe, ventral nerve cord, and [wing pouch + hinge] are shown. Genotypes and number of tissues measured for each genotype are indicated in parenthesis under the graphs. The x in the boxes indicates the mean value and the line inside the box indicates the median. Two samples t-tests for each genotype pair were performed using Microsoft Excel (Redmond, WA). When the p-value indicates that the pair variance is statistically significant, it is shown as a horizontal line across the genotypes with * (P<=0.05), ** (P<=0.01), *** (P<=0.001), or **** (P<=0.0001).

Given the potential complications associated with a phenotypic analysis of the *trus^1^* allele, we generated new *trus* null alleles using CRISPR/Cas9 targeted mutagenesis [47]. Two guide RNAs were designed to flank the entire exon2 and the following intron (Fig. 1A, magenta). We isolated and sequenced a total of 10 *trus* deletion lines (data not shown). Among the alleles, *trus^4-15^* has a single bp deletion at 3R:12,456,811 within target A. This results in a truncated 42 amino acid peptide with identity up to Arg40, two missense codons, and a premature stop (Fig. 1A). Since this peptide is so small even if it is stably expressed, we consider *trus^4-15^* to be a *trus* null allele. A second allele, *trus^35-2^* is a deletion of 1184 bp between 3R:12,456,805 and 3R:12,457,988 starting 3 bp upstream of target A and ending 5 bp downstream of target B. This deletion gives rise to a shorter 110 amino acid peptide lacking residues between Trp38 and Phe414 and (Fig. 1A).

Western blot analysis using a polyclonal antibody raised against full-length recombinant Trus showed a band migrating around 60 kDa, corresponding to the full-length Trus protein (predicted Trus molecular weight is 53.2 kDa), that is missing in *trus^4-15^/trus^4-15^*, *trus^35-2^/trus^35-2^*, and *trus^1^/trus^1^* larval extracts, while the protein level was reduced in *trus^4-15^*/TM6, *trus^35-2^*/TM6, and *trus^1^*/TM6 larval extracts (Fig. 1B). We detected no additional truncated fragments that reacted to the anti-Trus antibody on our Western blot (data not shown); however, it remains possible that either *trus^35-2^* and/or *trus^1^* could produce truncated fragments, since their phenotypes (described below) indicate that they behave as hypomorphs or antimorphs, respectively.

### *trus* mutants show developmental delay and pre-pupal lethality

To investigate phenotypes of the CRISPR/Cas9-induced *trus* mutants, *trus^4-15^*, *trus^35-2^*, and *Dftrus* chromosomes were balanced over TM6B *P[Dfd-GMR-nvYPF]Sb* and crossed in different combinations. First, we noticed that the *trus* mutant animals that were *Dfd-GMR-nvYFP* negative displayed higher embryonic lethality compared to *Dfd-GMR-nvYPF* positive animals. We determined the hatch rate of *trus^4-15^/Dftrus* embryos to be 35% (n=101), whereas the hatch rate of *Dfd-GMR-nvYPF* positive embryos was 88% (n=91) (Table 1). Those *trus* mutants that hatched exhibited extensive developmental delay and pupariated at 10-12 days AEL (After Egg Lay), which was 5-7 days later than *w^1118^* (Fig. 1C). *trus^4-15^/Dftrus* and *trus^35-2^/Dftrus* larvae did not actively crawl to a high position on the vial wall during the 3^rd^ instar stage, but instead stayed near the food surface. Some of the larvae remained in the wandering stage for up to 7 days, consistent with the phenotype observed with *trus^1^/Dftrus* mutants. After pupariation, most of the larvae died without becoming pupae. Noticeably, the pupariation rate of *trus^4-15^/ trus^35-2^* animals was 15% higher than *trus^4-15^/Dftrus* and *trus^35-2^/Dftrus* (Fig. 1C), and some of these pupated and developed to pharate adults, although none eclosed. Homozygous *trus^4-15^* larvae showed developmental delay and lethality equivalent to *trus^4-15^/Dftrus* (not shown). Homozygous *trus^35-2^* mutants showed developmental delay equivalent to *trus^35-2^/Dftrus*; however, the wandering stage larvae more actively wandered away from the food and some of them developed to pharate adults that did not eclose (not shown). Although, as described above, *trus^4-15^/Dftrus*, *trus^35-2^/Dftrus*, and *trus^4-15^/ trus^35-2^* produced significant embryonic lethality, those that hatched to become 1^st^ instar larvae largely survived through the larval stages and the pupariation rate of the different mutant allele combinations reached 70-85%, although only after substantial developmental delay. (Fig. 1C).

**Table 1.** *trus* mutant displays higher embryonic lethality.

| Marker | Genotype | plate # | # of embryos aligned | hatched embryos | % hatched |
| --- | --- | --- | --- | --- | --- |
| *Dfd-GMR-nvYFP-* | *trus*<sup>4-15</sup>/*Dftrus* | 1 | 41 | 14 | 35% |
|  | *trus*<sup>4-15</sup>/*Dftrus* | 2 | 60 | 21 | 35% |
|  |  |  |  | Average | 35% |
| *Dfd-GMR-nvYFP+* | *trus*<sup>4-15</sup>/*TM6B P[Dfd-GMR-nvYFP]Sb* or *Dftrus/TM6B P[Dfd-GMR-nvYFP]Sb* | 3 | 40 | 36 | 90% |
|  | *trus*<sup>4-15</sup>/*TM6B P[Dfd-GMR-nvYFP]Sb* or *Dftrus/TM6B P[Dfd-GMR-nvYFP]Sb* | 4 | 51 | 44 | 86% |
|  |  |  |  | Average | 88% |

A cage was set up for a cross between *trus*<sup>4-15</sup>/*TM6B P[Dfd-GMR-nvYFP]Sb* and *Dftrus/TM6B P[Dfd-GMR-nvYFP]Sb*. Embryos were collected on apple-juice/agar plates with yeast paste for 12 hrs. Embryos were sorted based on *Dfd-GMR-nvYFP* plus or minus, and separately transferred and aligned along a line of yeast paste on new apple juice/agar plates. The embryo hatch rates were counted 36-48 hrs. AEL.

The *trus^4-15^/Dftrus* trans-heterozygous combination is a protein null based on genomic sequencing and Western blotting results (Fig. 1B), and it produces consistent pre-pupal lethality and developmental delay. We used this genotype as representative of the ‘*trus* zygotic null mutant phenotype’ in most of our subsequent experiments.

### *trus* mutants show defects in tissue growth and cell proliferation

To determine what causes the developmental delay and lethality in *trus* mutants, we first dissected 3rd instar wandering larvae just before pupariation and performed immuno-fluorescent staining of imaginal discs and brains with anti-phospho-HistoneH3 (anti-PH3) antibody and 4’,6-diamidino-2-phenylindole (DAPI). Fig. 1D shows representative confocal images of brains (top two rows) and wing discs (bottom two rows) from wild type (Fig. 1D, a-d: *w^1118^*), *trus* mutants (Fig. 1D, e-m), and the *Zfrp8* mutant (Fig. 1D, n-q). We first noticed that in *trus^4-15^/trus^35-2^* and *trus^4-15^/Dftrus* larvae, brains were significantly smaller than wild-type, and the brain lobes looked unstructured, meaning there was no characteristic ring-structure in the optic lobes as normally appears in wild-type brain lobes (Fig. 1D, e and i, compared to a). Their ventral nerve cords (VNC) were narrower and elongated, and the surface of the entire brain appeared disrupted (Fig. 1D, e and i).

We also examined tissues from *Zfrp8^P^/DfZfrp8 (Df(2R)BSC356)* larvae to determine whether the phenotype of *Zfrp8* mutant larvae is similar to that of the *trus* mutant. *Zfrp8^P^* is a P-element insertion in the 5’-UTR of the *Zfrp8* gene, and it is not a protein null, therefore it is expected to show a mild phenotype [48]. *Df(2R)BSC356B* is a deficiency chromosome that lacks 42 genes including *Zfrp8*. Brains from *Zfrp8^P^/DfZfrp8 (Df(2R)BSC356)* larvae (Fig. 1D, n) were smaller than wild type (Fig. 1D, a) but not as small as *trus* mutants (Fig. 1D, e and i). The *Zfrp8* null allele *Df(SM)206* combined with the *Zfrp8^M-1-1^* allele that deletes part of the *Zfrp8* gene (*Df(SM)206/Zfrp8^M-1-1^*) is lethal during the early larval stage; therefore we were unable to obtain 3rd instar larval tissue to analyze [30].

We next examined the imaginal discs including leg, haltere, and wing discs from *trus^4-15^/Dftrus* larvae (Fig. 1D, k) and found that they were smaller than wild type (Fig. 1D, c), with wing discs showing the most severe growth impairment and morphological defects. The discs were under-developed and too small to dissect out from early wandering larvae; however, after additional development during the prolonged wandering stage the wing discs reached a size that could be dissected and further examined. Intriguingly, wing discs from *trus^4-15^/trus^35-2^* larvae developed inconsistently during the late wandering stage. Some were larger than wild type and often showed excessive folds, whereas the others were smaller than wild type, so the overall size distribution was diverse. Wing discs from *Zfrp8^P^/DfZfrp8* were less affected than *trus^4-15^/Dftrus* but were still smaller than wild type (Fig. 1D). Fig. 1F shows quantification of PH3 positive cells (number/mm^2^) and area (mm^2^) of brain lobe, ventral nerve cord, and wing pouch plus hinge as indicated with yellow-line enclosed areas in Fig. 1E. In brain lobes from *trus^4-15^/Dftrus* and *trus^4-15^/trus^35-2^* larvae, PH3 positive (mitotic) cell numbers were significantly less than *w^1118^* control, and the size (area) of brain lobes was also significantly smaller than *w^1118^*. We observed less significant changes of PH3 count of ventral nerve cords from *trus^4-15^/Dftrus* and *trus^4-15^/trus^35-2^*; however, size (area) reductions were still significant for the *trus* mutants comparing to wild type.

We saw significant decrease of PH3 positive cells in brain lobes from *Zfrp8^P^/DfZfrp8.* The quantification confirmed a significant reduction of mitotic cell number and area in wing pouch + hinge from *trus^4-15^/Dftrus* compared to wild type. Area reduction of wing pouch + hinge region from *Zfrp8^P^/DfZfrp8* mutants compared to wild type was also significant (Fig. 1F).

In contrast to the absolute pre-pupal lethality of *trus^4-15^/Dftrus* larvae, a small number of *trus^4-15^/trus^35-2^* larvae developed to pharates that did not eclose. The size and cell proliferation variances observed in wing discs from *trus^4-15^/trus^35-2^* animals seem to be consistent with the non-penetrate pre-pupal lethality of the mutant. We speculate that the *trus^35-2^* allele may produce a truncated Trus protein product that has partial function, and that may be enough to trigger late metamorphosis into the pupal-pharate stage.

### The core structure of Trus protein and its orthologs are conserved

Fig2A shows the domain structures of *Drosophila* Trus and Zfrp8 together with several vertebrate and yeast orthologs. Trus and Zfrp8 have highly conserved N-terminal (shown in green and light blue) and C-terminal (shown in magenta) globular domains. The C-terminal conserved domain was previously relegated to the PDCD2L /PDCD2 protein superfamily and named PDCD2_C in the Pfam protein database [49]. We call the N-terminal conserved domain PDCD2_N in this paper. AlphaFold prediction of the *Drosophila* Trus 3D structure is shown in Fig. 2B [50, 51]. Colored domains assigned in Fig. 2A are shown in the same color in Fig. 2B. Trus consists of a core module that includes two β-sheets facing each other (green and magenta), a pair of interacting β-strands (light blue and blue), and unstructured loops (shown in grey) (Fig. 2B). The Predicted Aligned Error (PAE) calculated by AlphaFold indicates high confidence in the relative position of scored residues 1-109 (PDCD2_N; shown in light blue and green) when aligned with residues 382-485 (PDCD2_C; shown in magenta) (Fig. 2C). This supports the packing between these regions that form a structural module despite the large unstructured loops (grey) that separate the PDCD2_N and PDCD2_C domains. The AlphaFold Trus structure indicates that the PDCD2_N consists of four β-strands from which three β-strands form a β-sheet structure immediately followed by an α-helix (shown in green). In addition, the first β-strand in the PDCD2_N domain (residues 9-16, shown in light blue) is predicted to form hydrogen bonds with another β-strand (residues 307-317, shown in blue) in the middle of the loop region between the domain PDCD2_N and the domain PDCD2_C (Fig. 2B). The alignment of the blue β−strands (residues 307-317) against both the N-terminal domain (PDCD2_N, residues 1-109) and the C-terminal domain (PDCD2_C, residues 382-485) shows high confidence based on the PAE, indicating packing of a core module that includes the N-terminal PDCD2_N (light blue and green, residues 1-109), the C-terminal PDCD2_C (magenta, residue 382-485), and a β-strand in-between (blue, residues 307-317) (Fig. 2C).

**Fig. 2.**
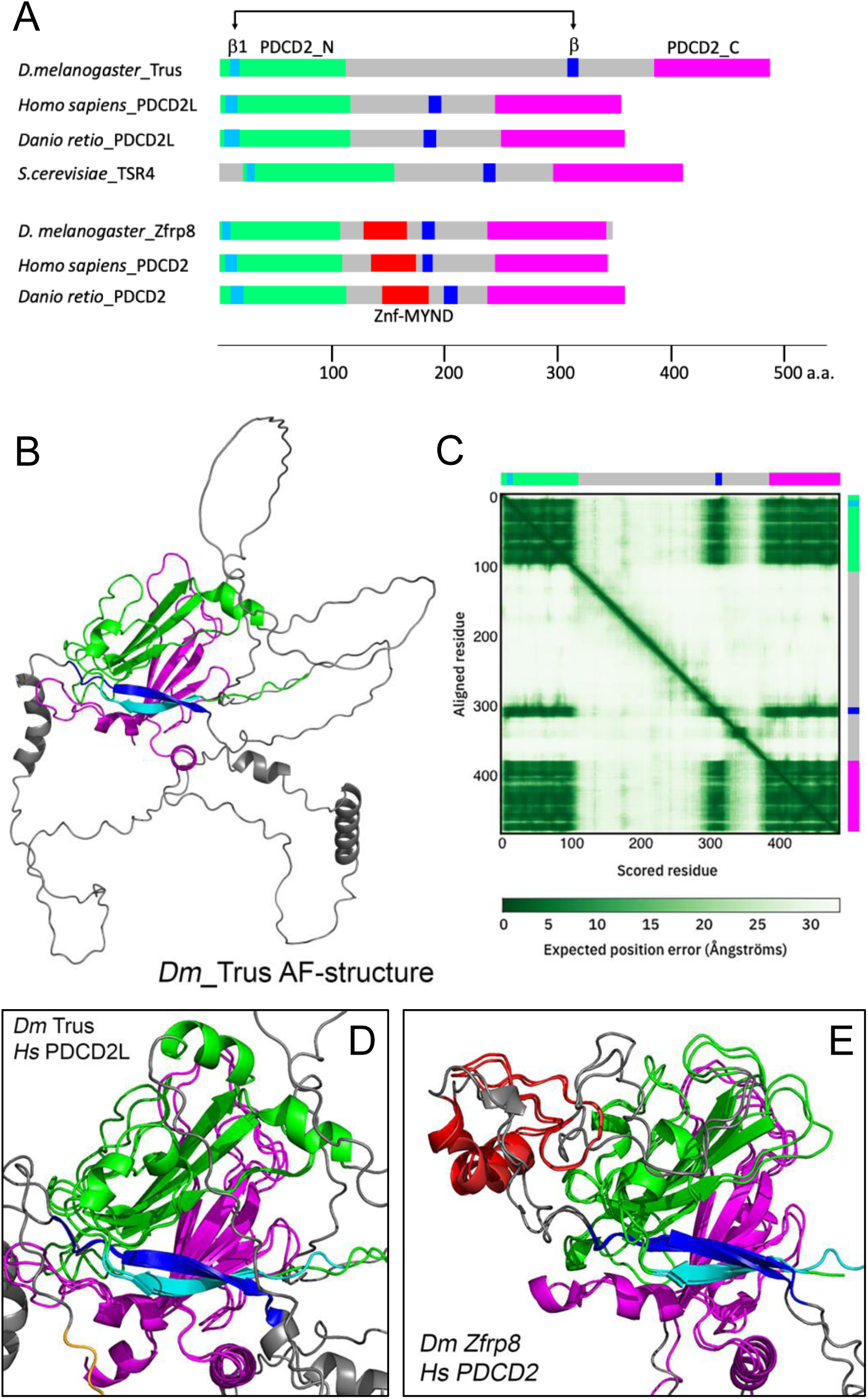
Predicted 3D Structure of Trus and its paralog Zfrp8 share a core module that is conserved through evolution. **(A)** Domain structure comparison of *Drosophila* Trus, its paralog Zfrp8, and their orthologs from different organisms. *D. melanogaster* Trus (Accession number: Q9VG62; Dmel\CG5333), Homo sapiens_PDCD2L (Q9BRP1), *Danio retio*_PDCD2L (Q5RGB3), *S. cerevisiae*_Tsr4 (P87156), *D. melanogaster*_Zfrp8 (Q9W1A3), *Homo sapiens*_PDCD2 (Q16342), and *Danio retio*_PDCD2 (Q1MTH6) are shown. (Green) PDCD2_N, (magenta) PDCD2_C, (red) MYND-type zinc finger domain (Znf-MYND), (light blue) first b-strand in PDCD2_N domain, (blue) b-strand that is predicted to interact with the first b-strand. (**B**) *D. melanogaster* Trus protein 3D structure predicted by AlphaFold (https://alphafold.ebi.ac.uk/). Color-coding of domains in the 3D structures are same as shown in A. **(C)** Predicted Aligned Error (PAE) of *Drosophila* Trus 3-D structure calculated by the AlphaFold. Color-coded bars representing the Trus protein shown in B are placed on upper and right sides of the panel. **(D)** Alignment of the core module of Dm Trus and Hs PDCD2L. **(E)** Alignment of the core module of Dm Zfrp9 and Hs PDCD2. 3-D structural alignments between orthologs were performed with PyMOL (Schrödinger LLC., NY).

Notably, structural alignments of *Drosophila* Trus against human PDCD2L (Fig. 2D), zebrafish PDCD2L (Fig. S3C), and *Saccharomyces cerevisiae* TSR4 (Fig. S3D) show striking conservations of the core module through evolution. *Drosophila* Zfrp8 is the ortholog to eukaryotic PDCD2 (<u>P</u>rogramme<u>d c</u>ell <u>d</u>eath protein <u>2</u>) [30]. Structural alignment of Zfrp8 against human PDCD2 indicates that the core structural module is also conserved through evolution in the Zfrp8/PDCD2 orthologs (Fig. 2E). We also note that AlphaFold predictions demonstrate that the core structural module is well aligned between Trus/PDCD2L and Zfrp8/PDCD2 paralogs (Fig. S3B), with the latter lacking the α-helix at the end of the PDCD2_N domain having instead a MYND-type zinc finger domain in the loop region outside of the core module that was previously shown to be involved in protein-protein interactions [52] (shown in red in Fig. 2E, Fig. S3A,B).

### Trus is expressed at high levels in larval mitotic tissues

We performed *in situ* hybridization with an anti-sense *trus* RNA probe to examine *trus* mRNA expression in 3rd instar wandering stage larval tissues. As shown in Fig. 3A, we detected *trus* expression at high levels in larval gut, ovary, brain lobe, wing disc, and lymph gland, where active cell proliferation is happening [53–56]. We also detected *trus* expression to a lesser extent in the salivary and prothoracic glands where cells are non-mitotic [57, 58].

**Fig. 3.**
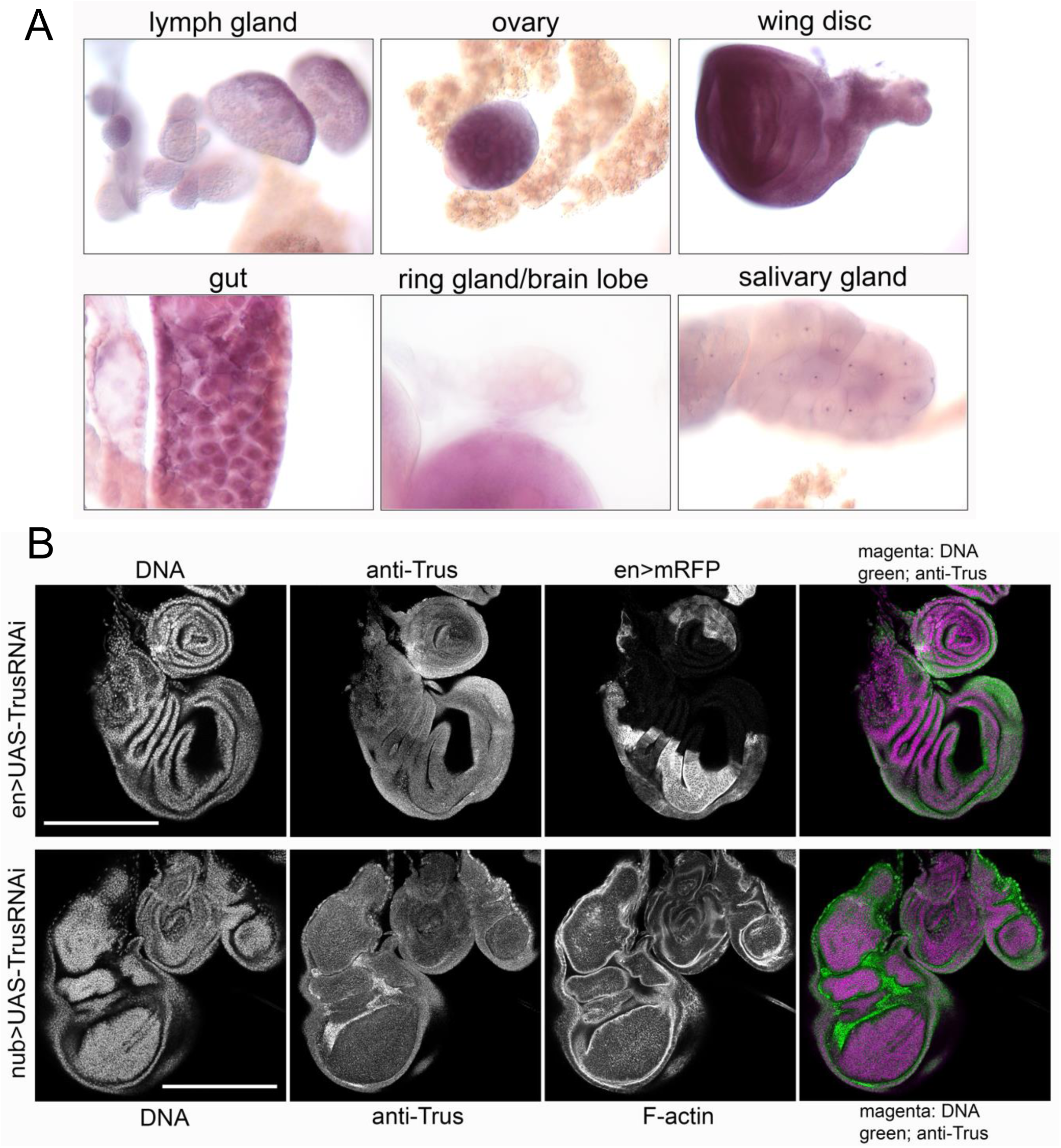
Trus expresses in mitotic tissues. **(A)** *In situ* hybridization with anti-sense RNA that hybridizes with *trus* mRNA reveals high level expression of *trus* in larval lymph gland, ovary, wing disc, gut, and brain lobe. Low expression is detected in ring and salivary glands. **(B)** anti-Trus antibody staining of tissues dissected from 3rd instar wandering larvae of *en-GAL4>UAS-TrusRNAi* or *nub-GAL4>UAS-TrusRNAi* larvae. Trus protein expression is detected in the entire wing, leg and haltere discs. The signals are reduced in the posterior half of the wing and leg discs (*en>UAS-TrusRNAi*) and in the pouch area of wing and haltere discs (*nub>UAS-TrusRNAi*), where *TrusRNAi* was induced.

To examine endogenous Trus protein expression and localization in larval tissues, we produced rabbit polyclonal antibody raised against full length recombinant Trus protein and affinity-purified the antibody using the antigen. As we described above, the affinity-purified anti-Trus antibody recognized the Trus protein which migrates around 60 kDa and is missing in either *trus^1^*, *trus^4-15^*, or *trus^35-2^* homozygous mutant larvae on Western blot (Fig. 1B). To reduce non-specific binding to larval tissue, the affinity-purified antibody was further pre-absorbed with fixed *trus^4-15^/Dftrus* larval tissues and used for immunostaining. We induced Trus RNAi in a part of tissue using either *en-GAL4* or *nub-GAL4* drivers and immuno-stained tissues from wandering 3rd instar larvae. In Fig. 3B, the upper panels represent wing and leg discs isolated from *en>TrusRNAi* larvae, showing Trus protein expression in the entire disc with reduced expression in the posterior half of the discs (Fig. 3B, anti-Trus), where *en-GAL4* induces mRFP expression (Fig. 3B, en>mRFP). The lower panels represent isolated wing, haltere, and leg discs from *nub>TrusRNAi* larvae, showing reduced Trus expression specifically in the pouch area of the wing and haltere discs where *nubbin* is expressed [59]. Taken together, these observations confirm that Trus protein is endogenously expressed in wing, leg, and haltere discs. Immunostaining with the affinity-purified, pre-absorbed anti-Trus antibody also revealed high level Trus protein expression in brain, leg and eye discs, and in the PG (Fig. S3).

### Trus localizes to the cytoplasm and is exported from the nucleus in a CRM1/XPO1-dependent manner

Immuno-detection of the endogenous Trus protein using anti-Trus antibody revealed that Trus localizes to the cytoplasm in larval tissue cells (Fig. S4). This is consistent with the observations of Landry-Voyer *et al.* [36] reported that PDCD2L, the human homolog of Trus, primarily localizes to the cytoplasm but shuttles between the nucleus and the cytoplasm, and the exportation from the nucleus is dependent on CRM1/XPO1, the RanGTP-binding exportin, which recognizes a leucine-rich nuclear export signal (NES) sequence. They showed that while PDCD2L primarily localizes to the cytoplasm, inhibiting CRM1 with Leptomycin B (LMB) or mutating the predicted NES sequence leads to retention of the protein in the nucleus [36]. We found that *Drosophila* Trus protein localizes in the same way as the human PDCD2L. As shown in Fig. S5A, EGFP-tagged Trus protein expressed in *Drosophila* S2 cells primarily localizes to the cytoplasm (No LMB); however, after Leptomycin B treatment, EGFP-Trus was retained in the nucleus, and longer LMB treatment was more effective (LMB 15min vs. LMB 115min), indicating that Trus shuttles between the nucleus and the cytoplasm in CRM1-dependent manner. We searched for a putative NES in Trus, based on NES consensus sequences reported in Kosugi *et al.*(2009) [60], and found at least 6 candidates in the loop region between the PDCD2_N and the PDCD2_C domains, which is much longer in Trus (273 a.a.) than human PDCD2L (132 a.a.) (shown in gray in Fig. 2A and B). Further investigations by mutating the candidates individually or in combinations are necessary to determine which NES candidate(s) is the Trus NES(s). We also over-expressed EGFP-tagged Trus *in vivo* using *UAS-EGFP-Trus* driven by *da-GAL4* and found that EGFP-Trus was ubiquitously expressed in larval tissues, and that the EGFP-Trus protein exclusively localized to the cytoplasm within the cell (Fig. S5B).

### Developmental delay of *trus* mutants is rescued by ecdysone feeding

20-hydroxyecdysone (20E) is the steroid hormone responsible for molting and metamorphosis in insects. We investigated if feeding 20E, or its precursor ecdysone, rescues the developmental delay of the *trus* mutant. The feeding scheme is outlined in Fig 4A. The larvae were fed mashed regular cornmeal fly food that was mixed with 20E or ecdysone dissolved in solvent (ethanol) beginning at the 1st instar stage. For the controls, fly food was mixed with either water or ethanol only. We found that feeding 20E to *w^1118^* larvae did not affect the pupariation timing (green in Fig. 4B) compared to the controls fed with water (blue) or ethanol (red), while feeding the precursor ecdysone slightly accelerated pupariation and reduced overall pupariation rate by 10% (purple) compared to the control or 20E-fed larvae (Fig. 4B). This data is consistent with the results reported previously by Ono (2014) where it was shown that feeding ecdysone to wild type larvae after L3 ecdysis accelerated pupariation by 6-12 hours and increased larval lethality [61].

**Fig. 4.**
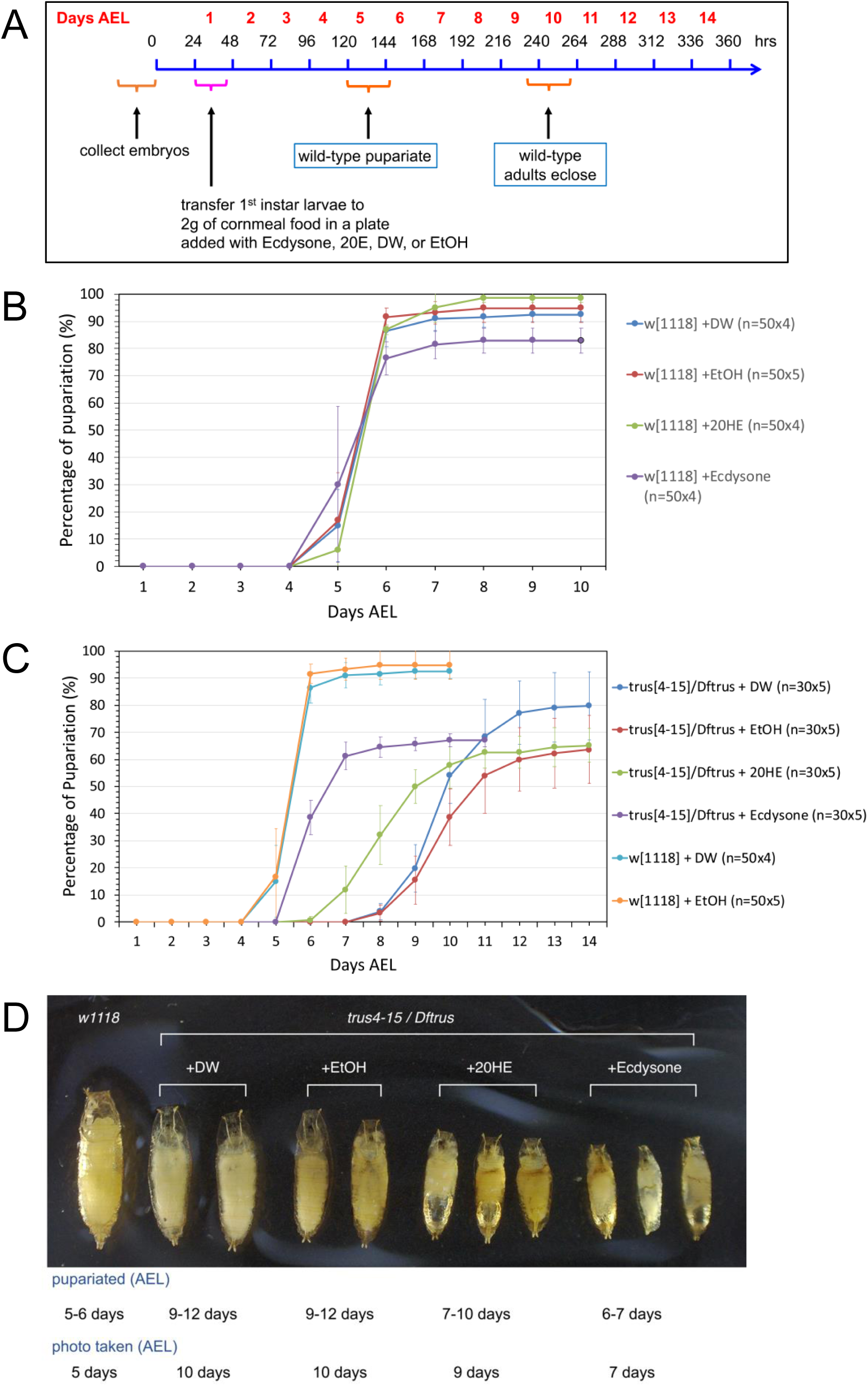
Ecdysone feeding to *trus* mutant larvae accelerates pupariation timing but causes precocious pupariation and does not rescue the pre-pupal lethality. (A) Schedule of the ecdysone-feeding experiment. **(B)** Pupariation timing of *w^1118^* larvae fed cornmeal fly food mixed with either ecdysone (purple), 20-hydroxyecdysone (20HE) (green), dH2O (blue), or ethanol (red). Each circle represents the average percentage of pupariated larvae from multiple dishes of the same genotype. Pupariated larvae were counted every 24 hours. Standard deviations are shown as vertical lines for each data point. **(C)** Pupariation timing of *trus^4-15^/Dftrus* larvae fed cornmeal food mixed with either ecdysone (purple), 20HE (green), dH2O (blue), or ethanol (red). Pupariation timing of *w^1118^*fed cornmeal food mixed with dH2O (light blue) or ethanol (orange) are shown as controls. **(D)** Pre-pupae that were fed cornmeal food mixed with either dH2O, ethanol, 20HE, or ecdysone with the time of pupariation indicated below. *w^1118^* pupa is shown on the left for size comparison.

When we examined *trus^4-15^/Dftrus* larvae fed with regular cornmeal food supplied with water (blue) or ethanol (red) they pupariated at 9-12 days AEL, while *w^1118^* larvae pupariated at 5-6 days AEL. 20E-fed larvae pupariated at 7-11 days AEL, and ecdysone-fed larvae pupariated at 6-7 days AEL (purple) (Fig. 4C). Interestingly, feeding ecdysone was much more effective in rescuing the developmental delay of *trus* mutants than feeding 20E, and as suggested by Ono (2014), this may indicate that ecdysone has an unknown function in regulating developmental timing separately from 20E.

Despite rescue of the developmental delay, 100% of the *trus^4-15^/Dftrus* animals that were fed 20E or ecdysone died as pre-pupae without developing into pupae. Notably, however those 20E-fed or ecdysone-fed pre-pupae were significantly smaller than the control mutant larvae (Fig. 4D) likely due to accelerated development, as has been noted previously in ecdysone feeding experiments [61].

### *TrusRNAi* in wing discs causes growth and cell proliferation defects

We next examined whether inhibiting Trus expression in wing discs affects tissue growth and larval developmental timing. *TrusRNAi* was induced either with *engrailed-GAL4* (*en-GAL4*), *en-GAL4* and *cubitus-interruptus-GAL4* (*ci-GAL4*) simultaneously, or *nubbin-GAL4* (*nub-GAL4*). Fig. 5A shows resulting wing phenotypes, in which *nub-GAL4* caused overall wing size reduction (*nub>UAS-TrusRNAi*), and *en-GAL4* (*en>UAS-TrusRNAi*) or *en-GAL4* and *ci-GAL4* (*en,ci >UAS-TrusRNAi*) caused disruption of wing veins and reduction of wing area, compared to the control (*UAS-TrusRNAi* without driver), especially in the posterior part of wings (Fig 5A). Quantification of the wing area using ImageJ indicates that the reduction in wing size is significant with either wing driver (Fig. 5B).

**Fig. 5.**
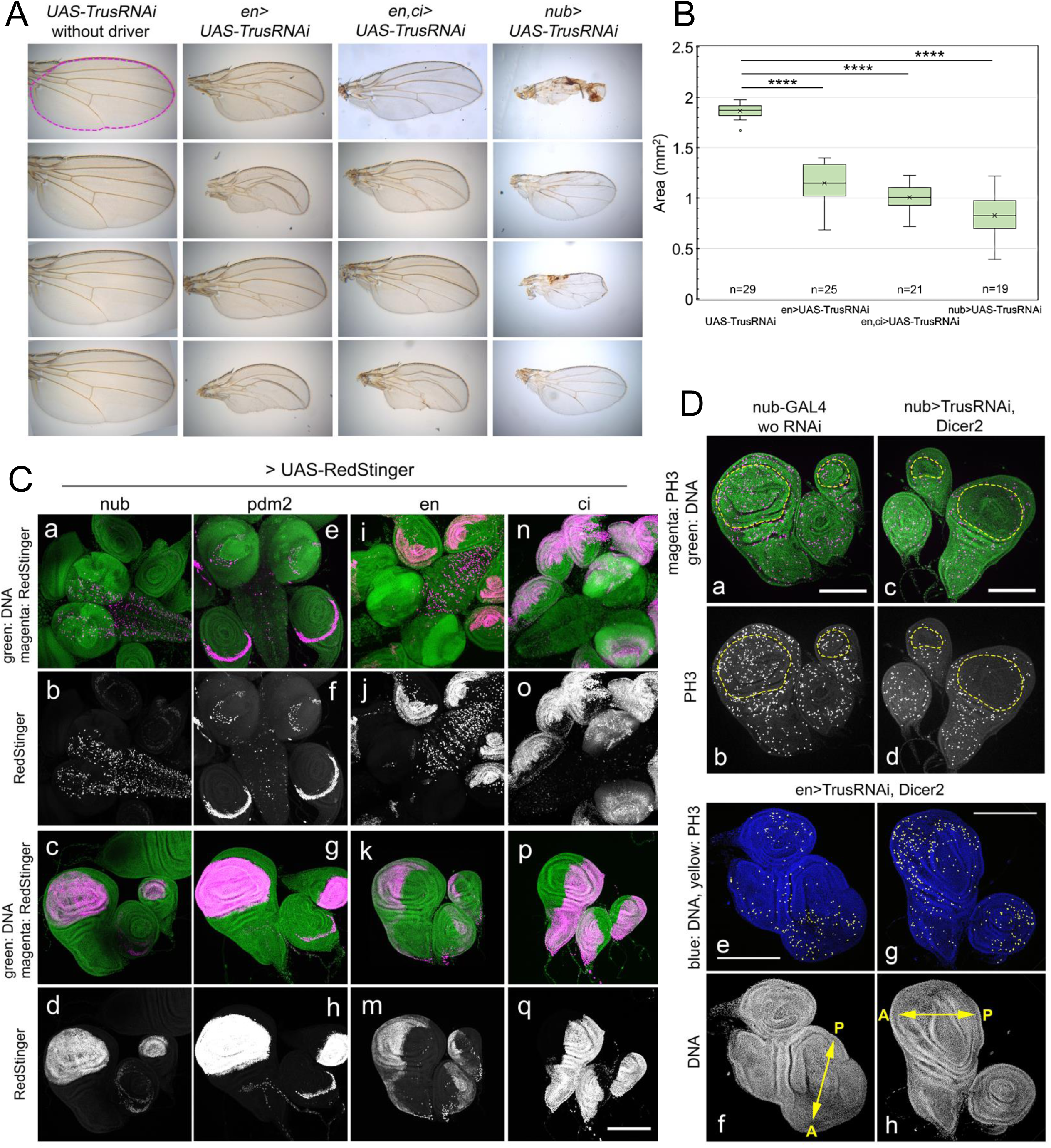
Trus RNAi induced with various wing disc drivers affects wing size and morphology and cellular proliferation. **(A)** Representative images of wings from female adult flies that had *TrusRNAi* induced with wing disc drivers. Adult wings from flies carrying *UAS-TrusRNAi* without any driver, *en-GAL4* driven *UAS-TrusRNAi*, *en* and *ci-GAL4* driven *UAS-TrusRNAi*, or *nub-GAL4* driven *UAS-TrusRNAi* are shown. **(B)** Quantification of wing area calculated in mm^2^. An example area is represented as magenta dotted outline in the top-left panel in A. ImageJ/Fiji (https://imagej.net) was used for area measurement. Box-and-whisker plots were generated, and two sample t-tests were performed using Microsoft Excel (Redmond, WA). The x in the boxes indicates the mean value and the line inside the box indicates the median. Sample numbers are indicated at the bottom of the graph (ex. n=29 for *UAS-TrusRNAi*). **** indicates P value <=0.0001. **(C)** *UAS-RedStinger* (shown in magenta) was induced with either *nub-GAL4* (a-d), *pdm2-GAL4* (e-h), *en-GAL4* (i-m) or *ci-GAL4* (n-q). DNA (DAPI, green) and RedStinger (magenta) staining are shown in first (a, e, i, n) and third (c, g, k, q) rows of images. RedStinger staining (white) is displayed in the second (b, f, j, o) and fourth (d, h, m, q) rows. Scale bar: 200μm. **(D)** Reduction of anti-PH3 stained foci is observed in the pouch area of wing and haltere discs in *nub>TrusRNAi, UAS-Dicer2* larvae (a-d) and the posterior half of wing discs and one half of the leg disc in *en>TrusRNAi, UAS-dicer2* larvae (e-h). In a and c, green: DAPI and magenta: anti-phospho-HistoneH3 (PH3) staining. Yellow dashed lines indicate wing and haltere pouch. In e and g, blue: DAPI and yellow: anti-PH3 staining. Arrows in f and h indicate the anterior-posterior axis of wing discs. Bar: 200μm.

Next we examined the expression pattern of each *GAL4* driver used in the *TrusRNAi* experiments, and later used in *trus* mutant rescue experiments (Fig. 5C). To analyze expression patterns in detail, we chose a fast-maturing nuclear reporter *UAS-RedStinger* (*DsRed.T4.NLS*) [62]. We confirmed that *nub-GAL4* and *pdm2 (POU domain protein 2)-GAL4* are both expressed at high levels in the pouch area of wing and haltere discs, as reported previously [59, 63] (Fig. 5C, c-d, g-h). In addition, *nub-GAL4* is expressed in some cells in the central brain at high levels and leg discs at low levels (Fig. 5C, a-b). On the other hand, *pdm2-GAL4* shows a distinct expression pattern in a restricted part of leg discs (Fig. 5C, e-f) and the posterior/ventral corner next to the pouch in wing/haltere discs (Fig 5C, g-h); *pdm2-GAL4* is not expressed in the brain except in lamina cells in optic lobe and a small number of cells in ventral nerve cord (Fig. 5C, e-f). *en-GAL4* and *ci-GAL4* are expressed in the posterior and anterior halves, respectively, in imaginal discs (Fig. 5C, i-m and n-q). In addition, *en-GAL4* is expressed in many cells of the VNC but not as much in the brain lobe (Fig. 5C, i-j), whereas *ci-GAL4* showed strong expression in the brain lobes including neuroepithelial cells that are differentiating as well as eye/antenna discs, but low expression in the VNC (Fig. 5C, n-o).

Further, we found that *TrusRNAi* caused a reduction of proliferation specific to the area where *TrusRNAi* was expressed. We observed that nub-GAL4 driven *TrusRNAi* leads to a reduction of mitotic cell number detected by anti-PH3 antibody only in the pouch of wing and haltere discs, resulting in shrinkage of the pouch (Fig. 5D, a-d,). en-GAL4 driven *TrusRNAi* reduced the mitotic cell number as detected by anti-PH3 antibody only in the posterior half of the wing disc and the leg disc, resulting in considerable shrinkage of the posterior half of the wing disc (Fig. 5D, e-h). Our observations indicate that *trus* disruption inhibits cellular proliferation cell-autonomously. We have shown in Fig. 5A that *TrusRNAi* driven by a combination of *en-GAL4* and *ci-GAL4* affects the posterior part of the wing more than the anterior (*en,ci>UAS-TrusRNAi*). This can be explained by the fact that when both *en-GAL4* and *ci-GAL* are used simultaneously, *en-GAL4* expression is much stronger than *ci-GAL4* for unknown reasons (data not shown).

To determine whether apoptosis is the cause of lethality and developmental delay in *trus* mutants, we examined whether apoptosis increased in *trus* mutant brains and wing discs. Using the TUNEL assay or anti-cleaved caspase3 antibody staining, we found no increase in apoptosis in *trus* mutant brains (data not shown). We detected some apoptosis in *trus* mutant wing discs; however, it was not consistently significant compared to wild type (data not shown). We further found that inhibiting apoptosis by ubiquitously overexpressing baculovirus p35, a caspase inhibitor [64, 65], with *daughterless-GAL4* did not rescue the developmental delay or lethality of *trus* mutants and did not rescue growth and cell proliferation defects of brain and wing discs (Fig. S2). Considering that brains and wing discs of *trus* mutants are smaller and have reduced mitotic cells compared to wild type, our results indicate that Trus is essential for tissue growth and developmental processes through its function in controlling cellular proliferation, but unlikely through apoptosis.

### *TrusRNAi* with wing disc drivers causes developmental delay and lethality

When we enhanced *TrusRNAi* using *UAS-Dicer2*, together with *nub>TrusRNAi* or *pdm2>TrusRNAi*, both of which express mainly in the wing pouch area (Fig. 5C, c-d, g-h), we noted a complete loss of wing blades leaving a small hinge in adult flies, haltere defects, and extra bristles (Fig. 6A). *Dicer2*-enhanced *nub>TrusRNAi*, *pdm2>TrusRNAi*, and *en>TrusRNAi* also resulted in lethality during pre-pupal or pupal stage. The pupariation rate stayed high with all the drivers; however, the adult eclosion rate of *nub>TrusRNAi* larvae decreased to 30% on average, and *pdm2>TrusRNAi* decreased to 10% on average, while the eclosion rate without a driver averaged 79% (Table2). Notably, *en>TrusRNAi* resulted in a high rate of embryonic lethality (not shown), and the surviving larvae were 100% pre-pupal lethal (Table2). In addition, many larvae showed a “Tubby-like” phenotype resulting in significantly smaller pre-pupae than *w^1118^* or *pdm2>TrusRNAi* pupae (Fig. 6B). This observation is in line with reports that *engrailed* mutant embryos are Tubby-like and lethal during the embryonic stage [66]. We suspect that *TrusRNAi* driven with *en-GAL4* causes inhibition of cell proliferation, specifically in posterior compartments, leading to a failure of posterior segment establishment during embryogenesis similar to *engrailed* mutants, and this may also contribute to the high rate of embryonic lethality.

**Fig. 6.**
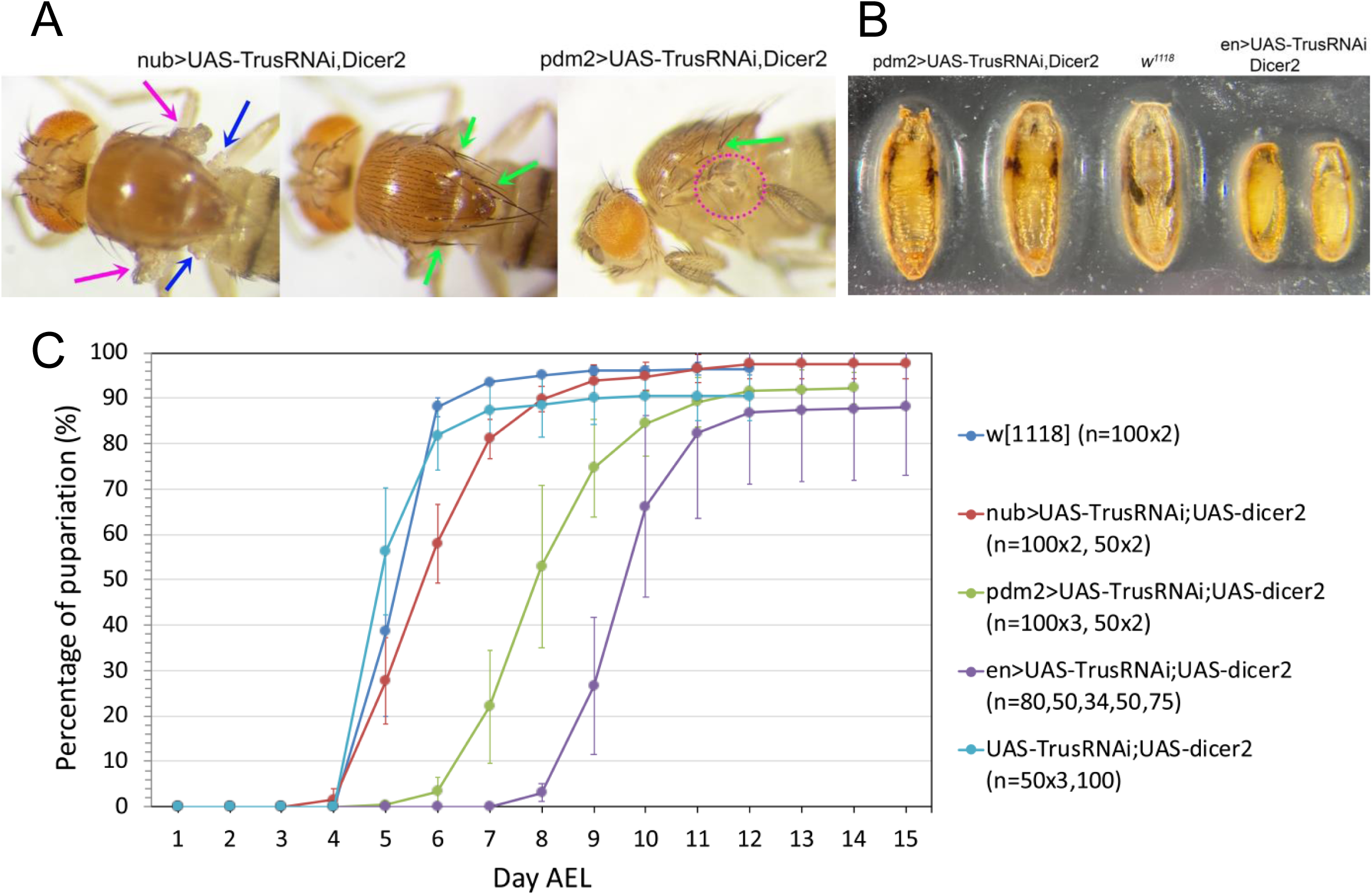
*TrusRNAi* induced with wing disc drivers delay pupariation timing. **(A)** *TrusRNAi* flies induced with either *nub-GAL4* or *pdm2-GAL4* in the presence of *UAS-Dicer2* show a complete loss of wing blade (magenta arrows), morphological defects in halteres (yellow arrows), and extra/disorganized bristles (green arrows). **(B)** (left) *pdm2>TrusRNAi* pupae are pharate with wing defects. (right) *en>TrusRNAi* larvae pupariate precociously resulting in smaller pre-pupae that never become pupae. (middle) *w^1118^* wild type pupa given for comparison. **(C)** Pupariation timing of *TrusRNAi* larvae induced with *nub-GAL4* (red), *pdm2-GAL4* (green), or *en-GAL4* (purple). Pupariation timing of *w^1118^* larvae (blue) and larvae carrying *UAS-TrusRNAi* and *UAS-Dicer2* alone (light blue) are shown as controls. Each data point represents an average pupariation percentage from multiple plates. The vertical line on each data point indicates the standard deviation. Numbers of 1st instar larvae picked at 1 Day AEL are shown in parentheses after the genotype.

*Dicer2*-enhanced *TrusRNAi* also caused significant larval developmental delay. Without a driver (*UAS-TrusRNAi*), pupariation occurred at 5-6 days AEL similarly to the *w^1118^* control (Fig. 6C light blue and blue). *nub>Trus RNAi* larvae pupariated at 5-8 days AEL (delayed 1-2 days compared to the control, Fig. 6C red), *pdm2>TrusRNAi* larvae pupariated at 7-10 days AEL (delayed 2-5 days, Fig. 6C green), and *en>TrusRNAi* pupariated at 9-11 days AEL (delayed 4-6 days, Fig. 6C purple). As we have shown above, *nub-GAL4* and *pdm2-GAL4* are expressed in the pouch area of wing and haltere discs (Fig. 5C), and *en-GAL4* is expressed at a high level in the posterior half of imaginal discs (Fig. 5C). Taken together with our results that *trus* mutant developmental delay was rescued by ecdysone feeding, we hypothesize that defects in growth and cell proliferation in wing/haltere/leg discs in *TrusRNAi* animals triggers a signal cascade leading to ecdysone deficiency and slow development.

### The Xrp1-Dilp8 stress response pathway is largely responsible for the developmental delay of *trus* mutants

The Xrp1-Dilp8 signaling pathway has been shown to coordinate *Drosophila* tissue growth with developmental timing [15, 67]. Xrp1 is a stress response transcription factor and is activated in growth impaired tissues promoting production and release of the peptide hormone Dilp8. Dilp8 then acts remotely on Lgr3 positive neurons in the central brain that inhibit the release of PTTH thereby blocking the production of molting hormone ecdysone [68]. Since *trus* mutants show growth and cell proliferation defects in imaginal discs, are developmentally delayed, and the delay is rescued by feeding ecdysone, we hypothesized that the Xrp1-Dilp8 pathway is the link between tissue growth impairment and developmental delay in *trus* mutants. To test this hypothesis, we recombined a genomic *GFP* insertion in *Dilp8* (*Dilp8-GFP*) [69] onto the *trus^4-^*^15^ chromosome. We found that *Dilp8-GFP* is expressed in wing, leg, and genitalia discs in *trus^4-15^/Dftrus* larvae, while the control larvae did not express *Dilp8-GFP* at all (Fig. 7A). As described previously, wing discs and, to some extent, leg discs dissected from these late wandering larvae are small and disorganized but show strong patches of *Dilp8-GFP* expression (Fig. 7B Dilp8-GFP). We also induced *TrusRNAi* with *nub-GAL4* and found that *Dilp8-GFP* was expressed in the pouch area of wing and haltere discs (magenta and yellow allows in Fig. 7C), suggesting that *Dilp8* expression was involved in the developmental delay observed with *TrusRNAi* described above (Fig. 6C).

**Fig. 7.**
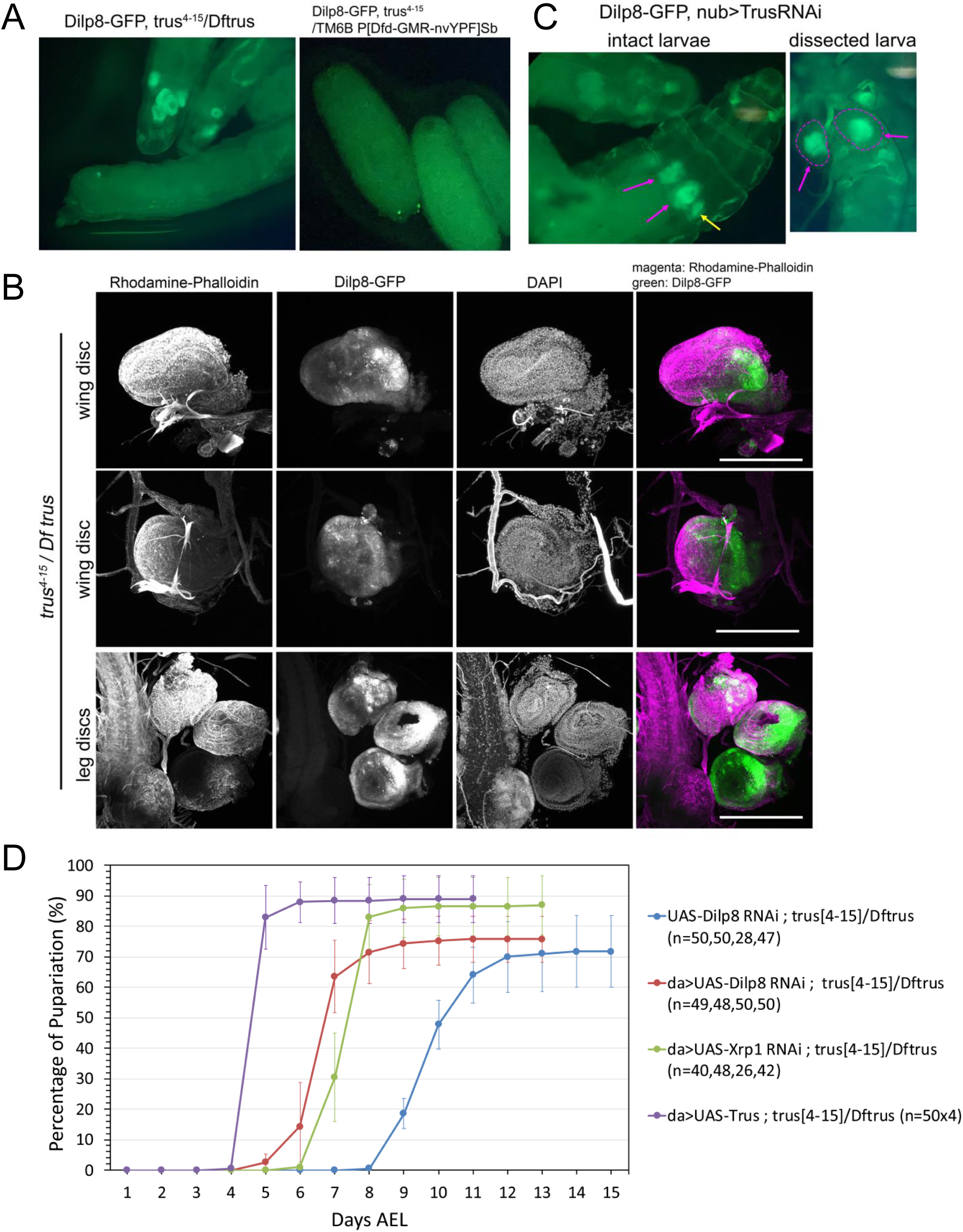
Xrp1-Dilp8 pathway is activated leading to developmental delay in *trus* mutants. **(A)** (left) Dilp8-GFP expression in 3^rd^ instar wandering *trus ^4-15^/ Dftrus* larvae. (right) Dilp8-GFP expression in 3rd instar wandering *trus^4-15^/ TM6B P[Dfd-GMR-nvYPF]Sb or Dftrus/ TM6B P[Dfd-GMR-nvYPF]Sb larvae.* Two bright GFP dots are Dfd-GMR-nvYFP signals on eyes. **(B)** Fixed and dissected wing (first and second rows) and leg (bottom row) discs from *trus ^4-15^/ Dftrus* 3^rd^ instar wandering larvae show Dilp8-GFP expression (green). Rhodamine-Phalloidin and DAPI stainings reveal significant reduction in size and abnormal morphologies of the discs. Scale bar: 200μm. **(C)** Expression of Dilp8-GFP in the pouch region of the wing disc (magenta arrows) and haltere disc (yellow arrow) from *nub> TrusRNAi* larvae. In the dissected larval image (right), wing discs are marked with a magenta dotted line and show GFP fluorescence in the middle area of the wing discs (wing pouch). **(D)** Developmental timing curves show that *da>Dilp8RNAi* (red) or *da>Xrp1RNAi* (green) significantly rescue the developmental delay of *trus^4-15^/Dftrus* larvae. Exogenous Trus expression from a *da>UAS-trus* transgene in *trus^4-15^/Dftrus* larvae rescues the developmental delay and lethality of the *trus* mutant (shown here in purple). *trus^4-15^/Dftrus* larvae with the *UAS-Dilp8RNAi* transgene alone serve as the negative control (blue).

To examine whether the Xrp1-Dilp8 signaling pathway is involved in the developmental delay of *trus* mutants, we individually knocked down *Xrp1* and *Dilp8* in *trus^4-15^/Dftrus* mutants by RNAi induced with *daughterless-GAL4*, a ubiquitous driver. We found that *Xrp1RNAi* and *Dilp8RNAi* accelerated the *trus* mutant’s pupariation timing by 4 and 5 days, respectively, compared to the control (Fig. 7D), indicating that the Xrp1-Dilp8 pathway is indeed responsible for much of the developmental delay.

### Identifying the tissues that require Trus protein activity during development

To determine which tissues require Trus function during development, we expressed Trus protein in *trus* mutant (*trus^4-15^/Dftrus*) animals from a *UAS-Trus* transgene using various *GAL4* drivers and examined whether *trus* developmental delay or lethality was rescued (Fig. 8A and Table2). Trus expression with a ubiquitously expressed *daughterless-GAL4 (da-GAL4)* rescued both developmental delay and lethality of *trus^4-15^/Dftrus* animals (Fig. 8A shown in blue). The eclosed adult flies had no defects and were fully fertile.

**Fig. 8.**
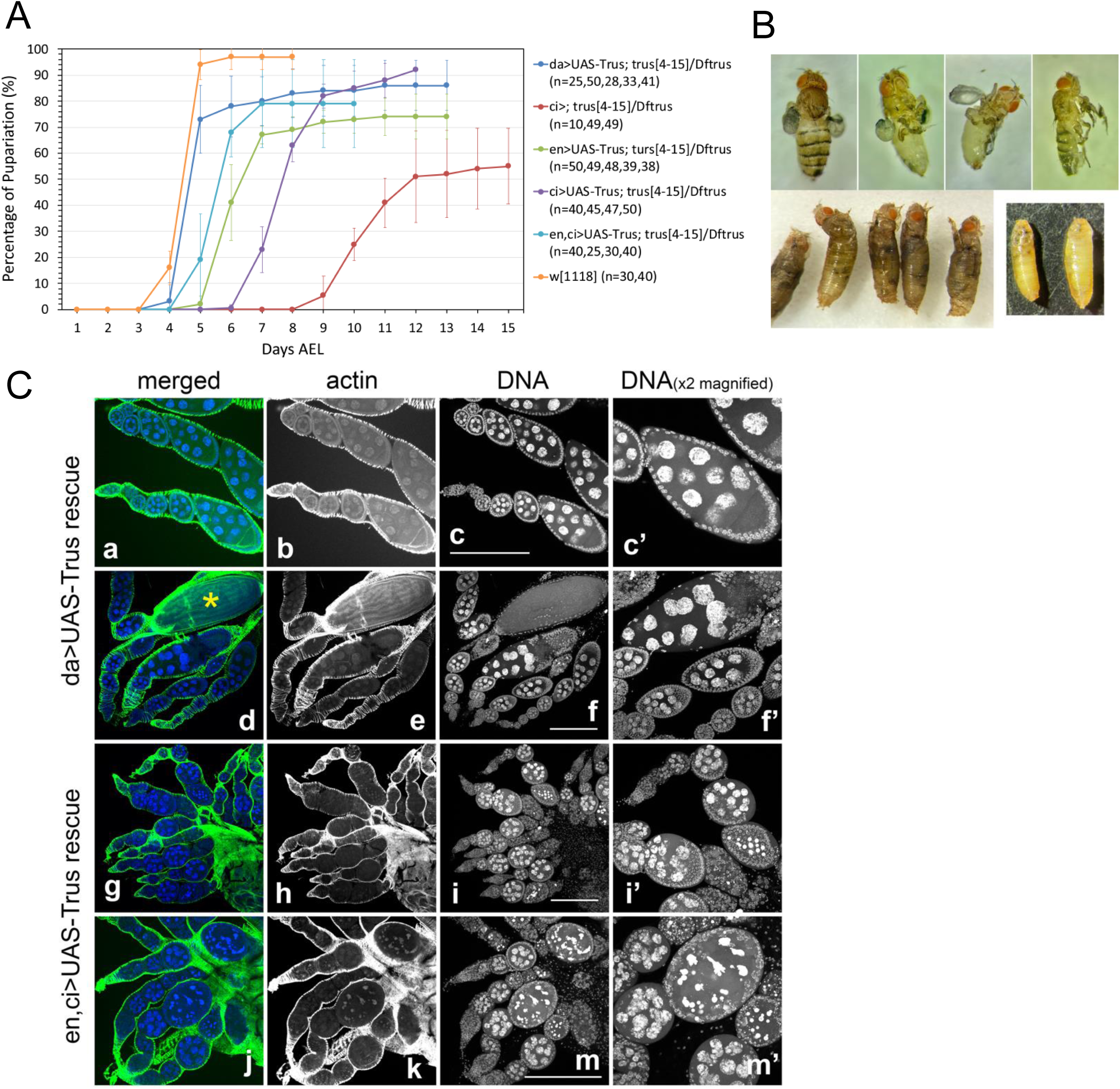
Rescue of lethality and developmental delay of the *trus* mutant can be achieved by Trus expression induced with specific drivers. (A) Developmental timing of *trus* mutants (*trus^4-15^/Dftrus*) with *trus* expression driven by various drivers display varying degrees of rescue. Compared to the *w^1118^* control (orange), *da-Gal4* (dark blue) rescues the best, with *en-Gal4,ci-Gal4* (aqua), *en-Gal4* (green), and *ci-Gal4* (purple) showing progressively lower degrees of rescue. *ci-Gal4 without UAS-Trus* (red) serves as the negative control. **(B)** Adult and pupal phenotypes of rescued lines. *ci>UAS-Trus* rescued *trus^4-15^/Dftrus* pre-pupal lethality with significant defects in wings, halteres, legs, and bristles (upper panels). Many flies eclose only half-way from the pupal case and die (lower left). *trus^4-15^/Dftrus* mutants without the *UAS-Trus* transgene arrest and die during the pre-pupal stage (lower right). **(C)** Ovariole phenotypes of rescued lines confirms that *da>UAS-Trus* rescue of *trus^4-15^/Dftrus* mutant females are fertile and show no defects in oogenesis (a-f, c’, f’). The yellow star indicates a mature egg produced (d). However, *en,ci>trus* in combination with *trus^4-15^/Dftrus* mutants, while being able to rescue mutant lethality, give rise to females that are sterile. Their ovarioles produce no mature eggs because egg chambers degrade at mid-oogenesis (g-m, i’, m’). (green) Rhodamine-Phalloidin (blue) DAPI. Scale bar: 200μm.

Trus expression driven by *ci-GAL4* or *en-GAL4* accelerated pupariation timing by up to 3 or 5 days, respectively, compared to the control, and a combination of *en-GAL4* and *ci-GAL4* accelerated pupariation timing by 6 days (Fig. 8A). Since *en-GAL4* and *ci-GAL4* alone express in a half of each imaginal disc complementing each other, rescue of developmental timing with Trus expression using both drivers may work in a synergetic way (Fig. 8A). We also found that *nub-GAL4* and *pdm2-GAL4*, which are strongly expressed in the wing pouch but not in a major part of the leg discs or the hinge/notum of wing discs, rescued neither the developmental delay nor lethality of the *trus* mutant (Table2). This is not surprising since Dilp8 is likely to be strongly expressed in the notum and hinge area of wing discs, leg discs, and other discs in *trus* mutants when Trus is expressed using *nub-GAL4* or *pdm2-GAL4*.

Importantly, while both *en-GAL4* and *ci-GAL4* driven Trus expression rescued the developmental delay of *trus* mutants, the lethality of *trus* mutants was rescued only by *ci-GAL4*. Most of the *trus* mutant larvae with *ci>UAS-Trus* pupated and reached the pharate stage. 18% of the animals eclosed or half-eclosed as adult flies with severe defects in legs and wings (Table2, Fig. 8B). As we have shown, expression of *en-GAL4* and *ci-GAL4* complement each other in wing and leg discs; however, their brain expression patterns are distinctly different. Notably, *ci-GAL4* is strongly expressed in a large number of cells in brain lobes including differentiating neural epithelia cells in optic lobes, whereas *en-GAL4* expression in brain is limited to some cells in the VNC (Fig. 5C). We speculate that perhaps it is this difference in central nervous system (CNS) expression of *ci-GAL4* vs *en-GAL4* that accounts for the rescue of *trus* lethality by one and not the other. The expression of *ci-GAL4* in CNS needs to be further investigated in detail in future studies.

### Trus is essential for oogenesis

In our rescue experiments, we expressed Trus using a combination of *en-GAL* and *ci-GAL4* together. This double driver further accelerated pupariation timing up to 6 days (light blue in Fig. 8A) and significantly rescued *trus* mutant lethality. 41% of 1st instar larvae eclosed as adult flies with no morphological defects (Table 2). However, we found that the eclosed females were completely sterile and laid no eggs. To determine when the block occurs in oogenesis, we dissected their ovaries and stained them with Rhodamine-Phalloidin and DAPI, As shown in Fig. 8C, in *en,ci>UAS-Trus* rescued ovaries, a normal number of egg chambers seemed to form, but nurse cell nuclei started to show abnormal morphologies at early stages and severe aggregation and fragmentation of the nuclear DNA by mid-oogenesis (stage 5/6) (Fig. 8C, g-m, i’, and m’). Additionally, follicle cells that normally surround the developing nurse cells and oocyte allow the egg chamber to hold its structure and change its shape from spheroid to elliptical as seen in the control (Fig. 8C, a-f, c’, and f’), also deteriorate and degrade by stage 5/6 in *en,ci>UAS-Trus* rescue (see Fig. 8C, i’ m’ comparing to c’ f’). Oogenesis seemed to arrest and egg chambers degraded completely after Stage5/6. Mature eggs always seen at the end of the egg chambers in the control ovaries (Fig. 8C, d, an egg shown with yellow star) did not exist in ovaries derived from *en,ci>UAS-Trus* rescued females. High-throughput transcriptome analyses have reported that both *engrailed* (*en*) and *cubitus interruptus* (*ci*) are not expressed or have very low expression in ovaries (http://flybase.org). Therefore, we infer that, although *en-GAL4* and *ci-GAL4* zygotically expressed *UAS-Trus* and rescued *trus* mutant (*trus^4-15^/Dftrus*) lethality, the drivers did not express *UAS-Trus* in ovaries, and thus did not rescue the Trus function that is essential for oogenesis.

**Table 2.** Percentage of pupariation and eclosion of larvae that were induced Trus RNAi in wing discs.

| Genotype<br>(number of 1st instar larvae) | number of pupae<br>or pre-pupae<br>(% of larvae) | number of adults<br>eclosed<br>(% of larvae) |
| --- | --- | --- |
| *w<sup>1118</sup>*-1 (n=100) | 95 (95%) | 95 (95%) |
| *w<sup>1118</sup>*-2 (100) | 98 (98%) | 94 (94%) |
| AVERAGE% | 97% | 95% |
| *nub>UAS-TrusRNAi,UAS-dicer2-1* (100) | 100 (100%) | 29 (29%) |
| *nub>UAS-TrusRNAi,UAS-dicer2-2* (100) | 100 (100%) | 33 (33%) |
| *nub>UAS-TrusRNAi,UAS-dicer2-3* (50) | 49 (96%) | 10 (20%) |
| *nub>UAS-TrusRNAi,UAS-dicer2-4* (50) | 47 (94%) | 19 (38%) |
| AVERAGE% | 98% | 30% |
| *pdm2>UAS-TrusRNAi,UAS-dicer2-1* (100) | 97 (97%) | 10 (10%) |
| *pdm2>UAS-TrusRNAi,UAS-dicer2-2* (100) | 86 (86%) | 11 (11%) |
| *pdm2>UAS-TrusRNAi,UAS-dicer2-3* (100) | 92 (92%) | 13 (13%) |
| *pdm2>UAS-TrusRNAi,UAS-dicer2-4* (50) | 47 (94%) | 4 (8%) |
| *pdm2>UAS-TrusRNAi,UAS-dicer2-5* (50) | 46 (92%) | 4 (8%) |
| AVERAGE% | 92% | 10% |
| *en>UAS-TrusRNAi,UAS-dicer2-1* (80) | 69 (86%) | 0% |
| *en>UAS-TrusRNAi,UAS-dicer2-2* (50) | 30 (60%) | 0% |
| *en>UAS-TrusRNAi,UAS-dicer2-3* (34) | 34 (100%) | 0% |
| *en>UAS-TrusRNAi,UAS-dicer2-4* (50) | 47 (94%) | 0% |
| *en>UAS-TrusRNAi,UAS-dicer2-5* (75) | 74 (99.7%) | 0% |
| AVERAGE% | 88% | 0% |
| *UAS-TrusRNAi;UAS-dicer2-1* (50) | 50 (100%) | 42 (84%) |
| *UAS-TrusRNAi;UAS-dicer2-2* (50) | 44 (88%) | 44 (88%) |
| *UAS-TrusRNAi;UAS-dicer2-3* (50) | 43 (86%) | 34 (68%) |
| *UAS-TrusRNAi;UAS-dicer2-4* (100) | 88 (88%) | 74 (74%) |
| AVERAGE% | 91% | 79% |

Number of 1<sup>st</sup> instar larvae indicated in the parentheses after genotype (ex. n=100) were selected and transferred to a new regular cornmeal agar plate. Number and percentage (in parentheses) of the larvae that became pre-pupae, pupae, and eclosed adults were counted and indicated in the second and the third columns.

**Table 2.** Rescue summary of *trus^4-15^/Dftrus* developmental timing and lethality.

| background | rescue by | pupariation<br>days AEL | days<br>rescued | lethality | places<br>express the GAL4 |
| --- | --- | --- | --- | --- | --- |
| *w*<sup>1118</sup> | n/a* | 5 | n/a | viable | n/a |
| *trus*<sup>4-15</sup>/*Dftrus* | n/a* | 12 | n/a | pre-pupal lethal | n/a |
| *trus*<sup>4-15</sup>/*Dftrus* | *da>UAS-Trus* ** | 5 | 7 | 89% eclosed<sup>#</sup><br>w/o defects, fertile | Ubiquitous [70] |
| *trus*<sup>4-15</sup>/*Dftrus* | *en, ci>UAS-Trus* ** | 6 | 6 | 41% eclosed<sup>#</sup><br>female sterile, oogenesis<br>stops at stage5/6 | *en + ci*<br>*en* expression is stronger than<br>*ci* when expressed together |
| *trus*<sup>4-15</sup>/*Dftrus* | *en>UAS-Trus* ** | 7 | 5 | pre-pupal lethal | posterior half of imaginal discs,<br>some cells in VNC **** |
| *trus*<sup>4-15</sup>/*Dftrus* | *ci>UAS-Trus* ** | 9 | 3 | 18% eclosed or half-eclosed<sup>#</sup><br>w/severe defects in<br>wings, halteres, legs | anterior half of imaginal discs,<br>neuroepithelial cell lineages in<br>optic lobes, low expression in<br>some cells in VNC **** |
| *trus*<sup>4-15</sup>/*Dftrus* | *nub>UAS-Trus* | 12 | 0 | pre-pupal lethal | wing/haltere disc (pouch),<br>some cells in central brain,<br>low expression in some cells in<br>leg discs **** |
| *trus*<sup>4-15</sup>/*Dftrus* | *pdm2>UAS-Trus* | 12 | 0 | pre-pupal lethal | wing/haltere disc (pouch<br>+ upper posterior corner of<br>hinge), a restricted area of leg<br>discs, lamina cells in optic<br>lobes, some cells in VNC **** |
| *trus*<sup>4-15</sup>/*Dftrus* | *repo>UAS-Trus* | 12 | 0 | pre-pupal lethal<br>rescued mobility in<br>wandering stage | pan-glial cells in brain [71] |
| *trus*<sup>4-15</sup>/*Dftrus* | *elav>UAS-Trus* | 12 | 0 | pre-pupal lethal | pan-neuronal cells in brain [71] |
| *trus*<sup>4-15</sup>/*Dftrus* | *e22c>UAS-Trus* | 12 | 0 | pre-pupal/pupal lethal<br>some reach to pharate<br>and die | epidermal cells[72], limited<br>cells in CNS (our observation) |
| *trus*<sup>4-15</sup>/*Dftrus* | *da>*<br>*UAS-Dilp8*<br>*RNAi* *** | 7 | 5 | pre-pupal lethal | ubiquitous |
| *trus*<sup>4-15</sup>/*Dftrus* | *da>*<br>*UAS-Xrp1*<br>*RNAi* *** | 8 | 4 | pre-pupal lethal | ubiquitous |

### Trus is found in a complex with SOP/RpS2 and eEF1a1

To search for possible Trus interactors, we used the Tap-tagging system described by Veraksa *et al.* (2005) [73], which identifies at least transiently stable protein complexes. *Drosophila* Kc cultured cells expressing a Tap-Trus fusion were lysed, and interacting proteins were isolated by affinity chromatography against the tag. The final eluate from the affinity column had two major bands and one band of somewhat lower intensity that were not found in controls (Fig. 9). One of these bands was (as expected) Trus itself; this band assignment was verified both by Western blotting with anti-Trus antibody and by mass spectrometry. It should be noted that the Trus band migrated on this gel around 70 kDa. Using mass-spectrometry, the other major band (band 3 on Fig. 9) is identified as String of pearls (Sop), the S2 subunit of the 40S ribosomal subunit (RpS2) [29], while band 2 is identified as eukaryotic translation Elongation Factor 1 alpha 1 (eEF1a1: CG8280), a factor that plays a role in shuttling tRNAs to the ribosome during translation [74]. To the best of our knowledge eEF1a1 has not been previously associated with either PDCD2L or PDCD2 containing complexes.

**Fig. 9.**
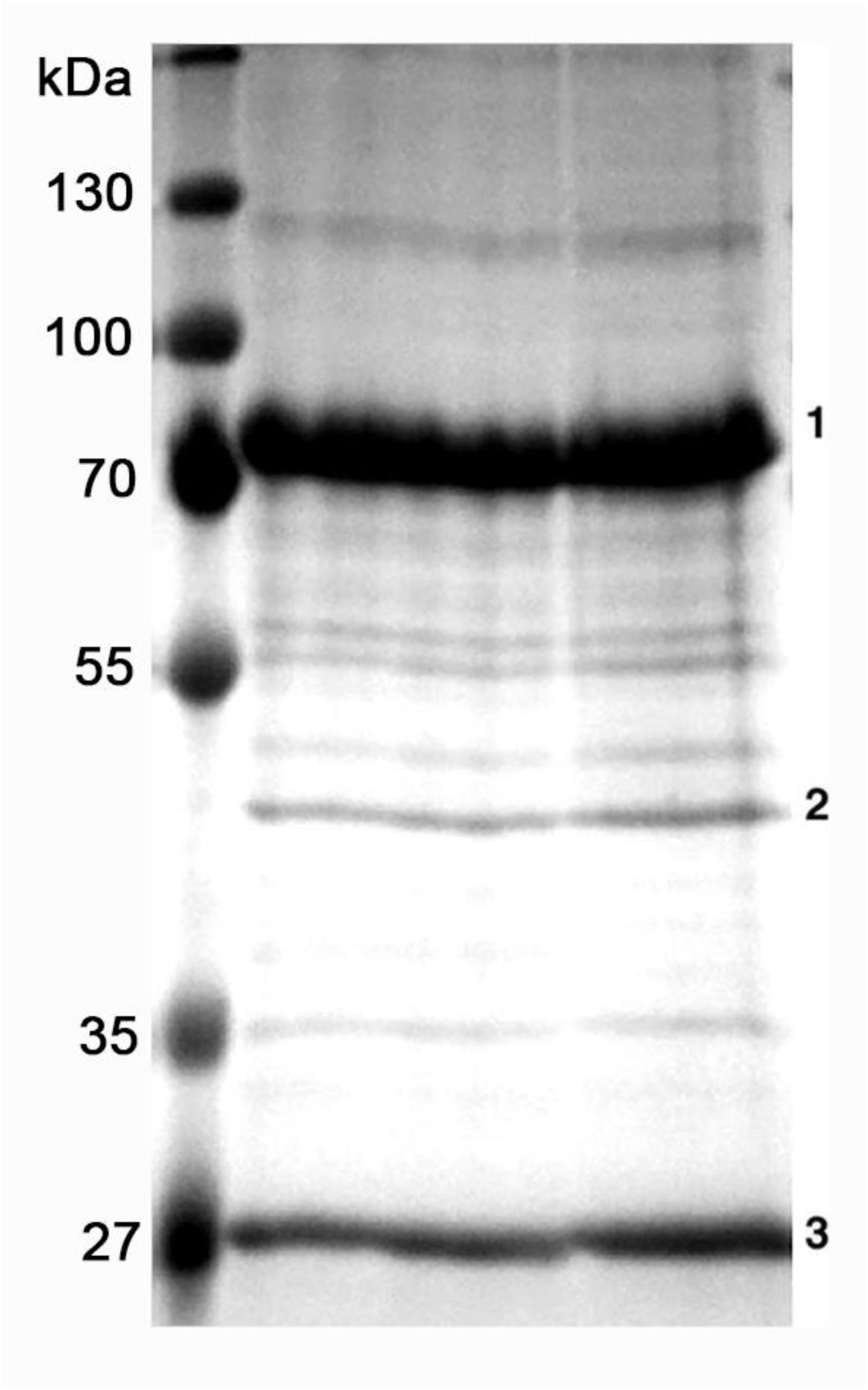
Tap-tagging reveals stable binding of Trus with Sop/RpS2 (String of pearls) and eEF1α1. Coomassie Blue staining of an SDS-PAGE gel after Tap-tagging with Trus. Three dominant bands are seen: (1) Trus (2) eEF1α1 and (3) Sop. IdentificaSons were made using MALDI-Mass spectrometry. To limit false posiSves, bands selected for mass spectrometry were unique when compared with proteins pulled down with a tag only construct (data not shown). Size marker in the left lane.

## Discussion

In this report, we characterized the phenotypes associated with mutations in the gene encoding Trus, the *Drosophila* homolog of the putative vertebrate ribosome subunit assembly factor PDCD2L. The most apparent phenotypes are observed in larval mitotic tissues such as imaginal discs and the central nervous system, while no obvious defects were noted in many endocycling tissue such as epidermis, fat body, muscle, much of the gut, and the prothoracic gland. One exception is the ovaries where endocycling nurse cells exhibit altered morphology at stage 6, just before the eggs degenerate. To a first approximation, these phenotypes suggest that dividing cells are most sensitive to loss of Trus, an inference consistent with the need for a large cellular ribosomal content for rapid growth. It is not clear what aspect of the cell cycle is affected in *trus* mutant larvae, but it is intriguing to note that *trus* was also uncovered in a transcriptome profiling screen of wing imaginal discs and S2 tissue culture cells as one of 63 genes that showed periodic transcription enriched at the G2 phase of the cell cycle in both cell types [75]. Furthermore, *RNAi*-mediated knockdown of *trus* in wing discs produced small abnormal adult wings, like what we report here, and cell cycle profiling of the *TrusRNAi* wing disc cells using fluorescence-activated cell sorting (FACS) showed increased G2/M [75], implying a role for Trus in cell cycle control at the G2/M transition.

A central question arises as to why a ribosome assembly factor would cause such a specific block in the cell cycle and not a more general global cellular defect. There are several potential explanations including the simple possibility that Trus is involved in multiple cellular processes. For example, vertebrate PDCD2L has been linked to apoptosis and has been found to be over expressed in many cancer cell lines, which could be consistent with Trus being a component of several different molecular machines, each affecting a specific cellular process [28, 40, 76–78]. Despite this caveat, the preponderance of data supports a primary role for PDCD2L/Trus and its paralogous couple, vertebrate PDCD2/*Drosophila* Zfrp8, in ribosome biogenesis [28]. PDCD2L appears to be the more ancient of the pair that duplicated prior to the divergence of animals, plants and lower eukaryotes to produce PDCD2 [38]. The core module of the PDCD2L/PDCD2 protein superfamily has been previously annotated as a TYPP domain by Burroughs and Aravind (2014). They identified the TYPP domain (named after <u>T</u>SR4, *<u>Y</u>wqG*, <u>P</u>DCD2L, and <u>P</u>DCD2) in yeast TSR4, bacterial DUF1963, and the PDCD2L/PDCD2 protein superfamily using extensive comparative genome sequence and structure analytical techniques. TSR4 is one of the proteins that were identified to be involved in rRNA processing in yeast *Saccharomyces cerevisiae* using large-scale genetic and computational screens [28, 38, 79]. The conservation of the core module among TSR4, PDCD2L and PDCD2 orthologs (Fig. 2, Fig. S3) suggests that these proteins exhibit similar biochemical properties, whereas the structural divergence between PDCD2L and PDCD2 paralogs, which includes the replacement of a particularα-helix with a MYND-type zinc finger domain (Fig. 2D,E, Fig. S3B), suggests that PDCD2L and PDCD2 may control specific functions or steps within the highly complex process of ribosomal biogenesis.

It was shown that PDCD2 co-translationally binds to newly synthesized uS5/RPS2, a stable component of the small (40S) ribosomal subunit, in both yeast and human cells, and was suggested to play a role as a dedicated chaperone for uS5/RPS2 to be incorporated into the pre-40S particle in the nucleolus [34, 35]. Supporting a direct connection of PDCD2L, a PDCD2 paralog, to ribosome biogenesis, PDCD2L also co-purifies with uS5/RPS2, and with PRMT3, a protein arginine methyltransferase [36]. Landry *et al.* (2016) further reported that, in contrast to PDCD2, human PDCD2L associates with several late 40S maturation factors and one of the last precursors of the mature 18S rRNA (18S-E pre-rRNA). Furthermore, they showed that PDCD2L has a functional NES and shuttles between nucleus and cytoplasm in a CRM1/XPO1-dependent manner [36]. In this report, we have shown that *Drosophila* Trus/PDCD2L also physically binds to *Drosophila* Sop/RpS2, and that it primarily localizes to the cytoplasm due to nuclear export since if the export machinery is blocked by Leptomycin B, an inhibitor of CRM1/XPO1, then Trus is predominately found in the nucleus. Taken together, current data suggests that PDCD2L plays a later role in the maturation process of pre-40S subunit than PDCD2, possibly acting as a protein adaptor for the CRM1/XPO1 in the pre-40S subunit nuclear exportation [28].

In many human cancer cell lines, PDCD2L appears not to be essential in contrast to PDCD2 or TSR4 genes [80] (https://depmap.org/portal/gene/PDCD2L), suggesting substantial functional redundancy between PDCD2L and PDCD2 or with other protein(s). However, in the context of normal development, PDCD2 and PDCD2L are both essential during embryonic stages with PDCD2L mouse knockout embryos being reabsorbed at mid-gestation while PDCD2 null embryos do not progress past the blastocyst/morula stages [39, 40]. In *Drosophila*, both Zfrp8/PDCD2 and Trus/PDCD2L are essential for cell proliferation, oogenesis, embryogenesis, and larval development [30, 42], suggesting that each paralog has a non-redundant function in each of these specific biological processes Our finding that Trus associates with SOP/uS5/RpS2, its cytoplasmic localization in the cell and *in vivo* tissues, and the ability to shuttle between nucleus and cytoplasm argue strongly that it is likely a functional homolog of vertebrate PDCD2L and that its primary role in *Drosophila* is also for small ribosome subunit maturation. In addition, certain aspects of the Trus loss-of-function phenotype similarly connect Trus function to ribosome biogenesis. Specifically, we find that *trus* hypomorphic alleles can produce rare escapers with phenotypes that include developmental delay, thin bristles and notched eyes which are the primary characteristics of *Minute* mutants. The *Minute* phenotype is produced by mutations in many ribosomal protein subunits. However, in the *Minute* case, the phenotype is observed in heterozygotes while with *trus,* heterozygotes are completely normal with respect to bristles, eye morphology and developmental timing. It is only when there is presumably less than 50% gene product that we see rare escapers with the classic *Minute* phenotype. We believe the likely explanation is simply that ribosomal subunits are required stoichiometrically as structural components of ribosomes, while Trus likely provides a temporal adaptor/transporter function to pre-40S subunits and can be recycled/shuttled back to the nucleus, making it less sensitive to heterozygosity.

Interestingly, the original *sop/uS5* mutants were also hypomorphs and showed classic *Minute* phenotypes [29]. It is also noteworthy that the *sop^P^* hypomorph is female sterile and blocks oogenesis at about stage 5 at which time nurse cell nuclear morphology becomes aberrant and degeneration occurs. This phenotype is strikingly similar to what we observe in *en,ci>UAS-Trus* rescued animals in which oogenesis is not rescued. However, we also observe that *en,ci>UAS-Trus* rescued egg chambers show severe deterioration of follicle epithelia by stage 5, which is earlier than *sop^P^* mutants (Fig. 8C) [29]. *Zfrp8* escapers are also reported to be female sterile [30]. Additional studies employing somatic and germline clones revealed a requirement of *Zfrp8* for proliferation of both germline stem cells and follicle stem cells. Germline clones progressed as far as stage 8 followed by degeneration [42], again similar to *sop* hypomorph alleles and the *en,ci>UAS-Trus* rescue egg chambers.

Given that Trus and Zfrp8 are paralogs and that their vertebrate counterparts appear to act as chaperone/adaptors during translocation of immature 40S components in and out of the nucleolus, it is perhaps not so surprising that they exhibit several common phenotypes, including lethality during all larval stages, developmental delay during the third instar, and oogenesis defects under hypomorphic conditions. Certain aspects of the null mutant phenotypes however are not identical. Minakhina *et al.* (2007) [30] noted that they obtained up to 5% escapers from *Zfrp8* null alleles while *trus* null larvae progress to prepupae, but none make it to adults. These differences, and the general polyphasic lethality we observe, could simply represent different degrees of perdurance of maternally supplied RNA or protein for the different genes. Another notable difference is that *zfrp8* loss-of-function alleles produce overproliferated lymph glands, perhaps due to an altered cell cycle [30]. In *trus* null mutants, the lymph gland is not particularly enlarged. However, the putative antimorphic/neomorphic *trus^1^* allele do exhibit enlarged lymph glands (Fig. S6). In this case, we envision that an N-terminally truncated Trus, that might arise from aberrant translational starts at downstream methionine codons, could act as a dominant negative form of Trus. Specifically, since PDCD2 and PDCDL2 appear to occupy the same binding site on uS5/RPS2, then an N-terminally truncated Trus might fully or partially block Zfrp8 activity leading to lymph gland overgrowth.

One other notable difference between Trus and Zfrp8 concerns their interaction partners. Mass spec analysis of proteins pulled down by an embryonically expressed N-terminally TAP tagged Zfrp8 identified 31 proteins including RPS2 (Tan et al.2016) which we also identified as an interacting partner of N-terminally TAP-tagged Trus. Interestingly, however, eEF1a1, which was a prominent band in our Trus pull down, was not identified as a component of Zfrp8 complexes. While this could result from the different tissues sources used for the pull downs, it is also consistent with the idea that each paralog has its own complement of unique interacting factors which then provides each complex with the potential for a distinct non-overlapping biological activity.

A key common phenotype of *trus, Zfrp8*, *M/+* and another ribosomal biogenesis component mutant known as *Noc1* is developmental delay during the third instar stage [4, 27, 30]. The cause of this phenotype has not been examined for Z*frp8* mutants but has been studied in *Minute* and *Noc1* mutants where it has been shown that subunit haploinsufficiency or improper ribosomal biogenesis triggers activation of the Xrp1 stress response pathway. In the case of *M/+* mutants, activation of the Xrp1 transcription factor requires input from RpS12, a ribosomal component, that appears to act as a sensor of ribosomal subunit concentration via an unknown mechanism, while in *Noc1,* the Eiger-JNK pathway is activated which can also induce Xrp1 [16, 27]. Once induced, Xrp1 can lead to a wide range of activities including blocks in translation and proteasome flux, activation of DNA repair and antioxidant genes, and, most relevant for this discussion, upregulation of *dilp8* in imaginal discs [8, 15, 26].

Dilp8 itself is a systemic stress signal that slows development by binding to a subset of Lgr3 positive neurons in the brain. These neurons synapse with the PG neurons that produce several developmental timing signals including PTTH, a neuropeptide that induces a rise in ecdysone production just prior to metamorphosis. Activation of the Lgr3 neurons by Dilp8 inhibits the rise in PTTH and thereby delays ecdysone production resulting in a pronounced developmental lag during the third instar stage [68, 81].

As we demonstrate in this report, Dilp8 expression is strongly activated by loss of Trus. Whole animal knockdown of either Xrp1 or Dilp8 ameliorates the developmental delay, as does feeding the larva ecdysone, pointing to this pathway as the prominent mechanism by which developmental delay is induced in *trus* mutants. Three additional points should be noted. First, the knockdown of either *Xrp1* or *Dilp8* does not totally rescue the developmental lag of *trus* mutants nor does feeding ecdysone. These animals still exhibit a 1.5 day delay relative to control larvae. While this could represent incomplete knockdown in the case of RNAi, or lack of obtaining the correct *in vivo* level of ecdysone in the feeding experiments, it is also possible that some other signal, perhaps from the brain, might also contribute to the delay phenotype and/or lethality. Second, we observe much better rescue of the delay phenotype when we use the molting hormone precursor α-ecdysone (E) as opposed to the actual molting hormone 20-hydroxyecdysone (20E). This difference in developmental timing when feeding E compared to 20E has been noted previously [61]. In that study, it was found that feeding E to wildtype larva early during the third instar stage, but not late, accelerated metamorphic timing and reduced overall body size compared to feeding 20E, which had only a minor effect on both parameters. These experiments suggest that E plays a distinct role during early third instar development that is divergent from the later role that 20E plays in triggering metamorphosis. It was suggested that E may help establish the larval minimal viable weight checkpoint. In our experiments, we fed E or 20E from the 1^st^ instar larval stage onward and noted that E accelerates metamorphic timing of *trus* mutants better than 20E which could indicate that *trus* larvae are primarily delayed during the early pre-minimal viable weight stage, and that is why they respond better to E rather than 20E in the feeding experiments.

A third consideration with respect to developmental delay is that *trus* larvae never make it past the pre-pupal stage, whereas *Ptth* mutants are viable as are *M/+* mutants, that also exhibit Dilp8-induced developmental delay. This suggests that there are likely additional hormone imbalances in *trus* mutants. One possibility is that high levels of Trus are needed in the prothoracic gland (PG) itself to enhance production of the 20E biosynthetic enzymes which are necessary for producing the large pre-metamorphosis inducing 20E peak. It is interesting to note that reduction in Noc1 has been shown to delay development both by inducing Dilp8 expression in imaginal discs and through direct effects in the prothoracic gland on E production [27]. In our case, however, RNAi knockdown of *trus* using PG-specific Gal4 drivers does not produce significant developmental delay on its own (data not shown).

Although the causative mechanism behind the developmental delay i.e. Xrp1-Dilp8 induction in *trus* mutants, is quite similar to that seen in *M/+* and *Noc1* mutants, once again there are some notable differences among the three. First, while we observe extensive proliferation defects in the CNS of *trus* mutants, such defects have not been reported for *M/+* or *Noc1* mutants. For *M/+*, this could simply be a dosage effect since the *trus* mutant is a homozygous null and may produce a more compromised cell than a *M/+* mutant which is a heterozygous knockdown of an individual RPS. For *Noc1,* null mutants were not examined and knockdown of *Noc1* in neurons did not produce a phenotype [27]. It would be interesting to determine if knockdown of an RPS or Noc1 in neuroblasts during larval stages produces a small brain phenotype like what we observe in *trus* mutants.

A second difference between *trus* and these other two types of alterations in ribosome biogenesis is induction of apoptosis through activation of cell competition. Heterozygosity for an RPS in *M/+* mutants results in pronounced apoptosis in imaginal discs and a simultaneous compensatory proliferation that allows the imaginal disc to reach its normal size and shape and to produce a normal appendage [22]. Inhibition of *M/+* induced apoptosis by expression of baculovirus p35 causes development of abnormal wing morphology indicating that apoptosis and the compensatory proliferation must be balanced to produce a normal structure. The apoptosis effect appears to be mediated to a large extend by Wg-induced cell competition [22]. RNAi knockdown of *Noc1* in wing discs also produces substantial apoptosis through both Eiger-JNK and Xrp1 induction [27]. In the case of *trus* mutants, we observe decreased, rather than increased, proliferation as seen in *M/+* mutants and no significant apoptosis within the discs or brain, as monitored by TUNEL assay or anti-cleaved caspase 3 staining. It seems that Xrp1 induction does not cause apoptosis in discs fully mutant for *trus* where there would not be any cell competition. Surprisingly, we also do not see apoptosis when we use *trus* RNAi to knockdown Trus in only a portion of the disc (i.e. *nub-GAL4>TrusRNAi*, *en-GAL4>TrusRNAi*). Additional studies will be required to address these differences.

In summary, our results highlight a role for the highly conserved ribosomal assembly protein Trus/PDCD2L in the control of mitotic proliferation, tissue growth, and oogenesis during *Drosophila* development. When the levels of this protein are reduced by mutation, the Xrp1-Dilp8 stress signaling pathway is triggered which slows development through inhibition of ecdysone production in an attempt to correct the cell proliferation and tissue growth imbalance.

## Materials and Methods

### Fly lines

Detailed genotype and sources of fly stocks that were used in this study are listed in Supplementary Table 1. Flies were reared on standard cornmeal fly medium at either 18, 25°C, or room temperature depending on the purpose.

### Production of *trus* CRISPR/Cas9 mutants

*trus* mutants were produced using the CRISPR/Cas9 system. Two target sequences in the *trus* gene (CG5333) that are unlikely to have off-target binding were identified using CRISPR Optimal Target Finder (http://targetfinder.flycrispr.neuro.brown.edu/) [82]. Target A is 5’-GGAATGGTCACCTCGTGTCTGGG-3’ and target B is 5’-GGATACGATCCCGCTGTTGGTGG-3’. Sense and antisense double strands of the targets A and B (see Supplementary Table 2) were cloned into *pU6-BbsI-chiRNA* (Addgene #45946) [47] for production of single stranded guide RNAs (sgRNAs) and were co-injected into vas-Cas9 embryos (BDSC_51323) by BestGene Inc. (Chino Hills CA). Single G0 adults were crossed to *w^1118^* and then 10 F1 males were crossed to *w^-^; Bl/CyOGFP; TM2/TM6B P[Dfd-GMR-nvYPF]SbTb.* From the established lines balanced over *TM6B P[Dfd-GMR-nvYPF]SbTb*, homozygous lethal lines were selected as *trus* null mutant candidates. Genomic DNA from balanced adults or homozygous larvae from each candidate line was prepared [83] and a 1.7kb *trus* genomic region was amplified by PCR (Expand High Fidelity PCR System, Millipore Sigma, Burlington, MA) with two primers, Trus2_for and Trus1_rev (Supplementary Table 2). Of the 5 pupal lethal lines from G0 #35, 4 had the same deletion of 1184 bp. Line 35-2 was selected and is hereafter called *trus^35-2^.* Of the 5 pupal lethal lines from G0 #4, 2 had the same 1 bp deletion at target A and were designed *trus^4-15^*. One line had a deletion differing in size from *trus^35-2^* but was not selected for further study. We selected line *trus^4-15^* and designated it *trus^4-15^.*Genomic DNA from *trus^1^/ TM6B P[Dfd-GMR-nvYPF]SbTb* adults was also prepared and sequenced as described above with Trus1_for, Trus3_rev, TrusB_for, and TrusC_for primers.

### Trus antibody production

Full length Trus protein N-terminally tagged with 6xHis-EGFP (His-EGFP-Trus) was expressed in Sf9 cultured insect cells using the Bac-to-Bac Baculovirus Expression System (Thermo Fisher Scientific, Waltham MA). The trus coding sequence, codon-optimized for insect cell expression and N-terminally tagged with 6xHis-EGFP, was cloned into pFastBac1 (pFB-His-EGFP-HRV3C - Trus) using NEBuilder High-Fidelity DNA Assembly (New England Biolabs, Ipswich MA). A HRV3C cleavage target sequence (3C) was inserted between EGFP and Trus. The plasmid was transformed into DH10Bac E. coli. cells and recombinant bacmids were isolated. Sf9 cells were seeded in a 6-well plate and transfected with the His-EGFP-HRV3C-Trus bacmid using CellFectinII (Thermo Fischer Scientific). The transfected cells were cultured at 28°C for 72 hrs. The supernatant (P0 virus stock) was collected and subsequently used to infect 10mL Sf9 cells to produce a P1 virus stock; the P1 was used to produce a 100mL P2 virus stock, and the P2 was used to produce a 300mL P3 virus stock. Virus infection of each amplification stage was confirmed under an optical microscope and His-EGFP-3C-Trus protein production was monitored by EGFP expression under a fluorescent microscope. Virus titer was measured for the P3 virus stock by Expression Systems (Davis CA) and used for subsequent His-EGFP-Trus protein expression. His-EGFP-3C-Trus recombinant protein was expressed in 2L of Sf9 cells, and the cell lysate was applied to a TALON CellThru metal affinity resin column (Takada Bio USA Inc., San Jose CA). The column was washed, and His-EGFP-3C-Trus protein was eluted with 150mM imidazole. The protein elution was monitored with EGFP fluorescence, and EGFP-positive fractions were collected and combined. The combined elution was incubated with HisGST-HRV3C protease to cut the His-EGFP tag at the 3C site, concentrated using Amicon Ultra-15 (MilliporeSigma, Burlington, MA), and incubated overnight at 4°C. It was then buffer-exchanged to lower imidazole concentration to 10mM using a Sephadex G-25/PD-10 desalting column (Cytiva, Marlborough MA) and combined with HisPur Ni-NTA resin (Thermo Fisher Scientific) and incubated at 4°C for 3 hours. The protein-resin solution was centrifuged, and the supernatant that contained Trus protein was collected, concentrated with Amicon Ultra-15, and loaded onto a Superdex-200 gel filtration column (MilliporeSigma). Trus protein containing fractions were combined, buffer-exchanged to Phosphate Buffer Saline (PBS, pH7.4) and concentrated. This concentrated purified Trus fraction was used to inject two rabbits to produce polyclonal anti-Trus antibody by Thermo Fisher Scientific. Anti-Trus antibody (AB2866) was affinity-purified from serum obtained from one of the rabbits with bacterially expressed recombinant Trus protein using AminoLink Plus Coupling Resin (Thermo Fisher Scientific) and was used for Western blotting (WB) at 1:2000. The affinity-purified anti-Trus antibody was diluted at 1:2000 and pre-absorbed against fixed *trus^4-15^/Dftrus* 3rd instar larval tissues and used for immuno-fluorescence (IF).

### Other antibodies and cell staining reagents

Phospho-histone H3 (Ser10) antibody (Cell Signaling Technology, Danvers MA) was used for IF at 1:1000 and a-Tubulin antibody (DM1A) (MilliporeSigma, T9026) was used for WB at 1:5000. Alexa 488-Goat anti-Rabbit IgG (H+L) and Alexa 555-Goat anti-Mouse IgG (H+L) Secondary Antibodies were used at 1:500-1:1000 for IF (Cell Signaling Technology). DyLight 680-Goat anti-Mouse IgG (H+L) and DyLight 800-Goat anti-Rabbit IgG(H+L) Secondary Antibodies were used at 1:10000 for WB (Thermo Fisher Scientific). For DNA staining, 4’,6-diamidino-2-phenylindole (DAPI) was used at 1.25mg/ml (MilliporeSigma), for F-actin staining, Rhodamine-Phalloidin was used (Thermo Fisher Scientific).

### Western blotting

Wandering 3rd instar larvae of *w^1118^* and *trus* mutants were collected and homogenized in SDS-PAGE sample buffer, boiled, and centrifuged to remove debris. Samples were run on 4-12% NuPAGE Bis-Tris Mini Protein gels (Thermo Fisher Scientific) together with the purified recombinant Trus protein fraction and the separated proteins were transferred to Polyvinylidene fluoride (PVDF) membrane. The membrane was blocked with x1/10 1xPBS1%Casein Blocker (BIO-RAD, Hercules CA), probed with purified rabbit anti-Trus antibody (this study) and mouse anti-α-Tubulin antibody (MilliporeSigma, T9026), washed, and then detected with DyLight secondary antibodies (Thermo Fisher Scientific). Finally, the membrane was washed and scanned with the Odyssey M imaging system (LICORbio, Lincoln NE).

### Immuno-fluorescence staining, imaging, and image analysis

Third instar wandering stage larvae were roughly, turned inside-out and removed most of the gut and the fat tissues removed in PBS, fixed with 3.2% EM grade Paraformaldehyde (Electron Microscopy Sciences, Hatfield PA) in PBS for 25 minutes, washed multiple times with PBST (0.3% Triton-X100, 1x PBS) on rotary shaker at room temperature. They were then incubated with primary antibody solution overnight at 4°C, washed with PBST and incubated with Alexa Fluor^TM^ conjugated secondary antibody solution (Thermo Fisher Scientific, Waltham MA) overnight at 4°C. Next day, they were rinsed with PBST, incubated with DAPI and/or Rhodamine-phalloidin for 5min. at RT, washed again multiple times with PBST. Brain and imaginal discs were dissected and mounted on a slide glass with anti-fading 1,4-Diazabicyclo[2.2.2]octane (DABCO) containing glycerin/PBS solution. Images were acquired with a Zeiss LSM 710 laser scanning confocal microscope using Zen imaging software (Zeiss, Jena Germany). To scan *Drosophila* tissues, 20x Plan-Apochromat/0.8 objective was used. An Argon laser, a HeNe laser, or a 405nm Diode laser were used to detect Alexa Fluor^TM^ 488 and EGFP, Alexa Fluor^TM^ 555 and mRFP, or DAPI, respectively. Normally, multiple z-series with 1.20μm interval covering the entire tissue thickness (pressed to 25-45μm) were scanned. For further image analysis including obtaining maximum projections and quantification of anti-PH3 foci count and area measurement, ImageJ/Fiji [84](https://imagej.net/software/fiji/) was used. Based on the image analyses, box-and-whisker plots were generated and two samples student t-test for each genotype pair was performed using Microsoft Excel (Microsoft, Redmond WA).

### *In situ* hybridization

To make the trus antisense *in situ* probe, the pFLC-1-trus cDNA clone (BDGP RE69372) was linearized with Eco53K1 and transcribed using DIG RNA Labeling Mix with T3 RNA polymerase (Roche, Indianapolis IN) according to manufacturer’s instructions. Wandering larvae of *w^1118^* and *trus^1^* were collected, dissected, and fixed. Hybridization using the digoxigenin-labeled antisense *trus* RNA probe and detection were performed as described previously [85, 86].

### Generating UAS-Trus and UAS-EGFP-Trus transgenics

To generate UAS-Trus and UAS-EGFP-Trus transgenics, the Trus coding sequence from an EST plasmid (Berkeley Drosophila Genome Project: RE69372) [87] and EGFP coding sequence from pEGFP-c1 (Clontech) were cloned into *pUAST.attB* [88] using NEBuilder HiFi DNA Assembly (New England Biolabs) and injected into embryos derived from a cross between flies with the *attP* recombination site inserted at 2L-VK00037 (BDSC_9752) and the *dPhiC31* integrase source (PhiC31 integrase source; BDSC_40161) for phiC31-mediated site-specific recombination (Bischof et al., 2007) and screened for *w+* transgenics that integrated *UAS-Trus* or *UAS-EGFP-Trus* (BestGene Inc., Chino Hills CA).

### Developmental Timing Assay

*trus^4-15^*, *trus^35-2^*, and *Dftrus* chromosomes were balanced over TM6B *P[Dfd-GMR-nvYPF]Sb*. As the *Dfd-GMR-nvYPF* fluorescent marker shows distinctive eye expression from embryonic stage 13 to adulthood, it is easy to score at the early larval stage [89]. These flies were crossed in different combinations. Embryos were collected on apple juice plates for 12-24 hours. 30-50 1st instar larvae (24-36 hours AEL) were screened for *Dfd-GMR-nvYFP* negative under a fluorescent dissecting microscope to obtain 1^st^ instar larvae of *trus^4-15^/Dftrus, trus^35-2^/Dftrus*, *trus^4-15^/ trus^35-2^*, *trus^4-15^* homozygous, or *trus^35-2^* homozygous and transferred to a standard cornmeal-agar fly medium in a 60mm culture dish. Another 60mm culture dish with a 20mm hole in the center covered with nylon mesh was placed as a lid and tightly taped together. 3-4 plates (total of at least 150 larvae) for each genotype were prepared and placed in humidified chambers at 25°C. Each plate was observed under a dissecting microscope throughout the subsequent development, and pupariation, pupation, and adult eclosion were assessed once every 24 hours for 10-18 days.

### Ecdysone Feeding Assay

α-Ecdysone (MilliporeSigma E9004) and 20-Hydroxyecdysone (MilliporeSigma H5142) were dissolved in ethanol at 10mg/mL and used as stock solutions. 50μL of either (1) water, (2) ethanol, (3) α-Ecdysone stock solution, or (4) 20-Hydroxyecdysone stock solution were mixed with 2g of mashed standard cornmeal/agar fly medium and placed in the center of 60mm plastic culture dish. The supplemented fly medium was air-dried for 2-3 hours to allow ethanol to evaporate. Cages were set up for *w^1118^* and a cross of *trus^4-15^/TM6SbTbHuYFP* x *Dftrus/TM6SbTbHuYFP,* and embryos were collected on apple juice agar plates for 12-24 hours. 50 1^st^ instar larvae of *w^1118^* or *trus* mutant (nvYFP negative) were transferred to the prepared supplemented or control media (1)-(4) for following developmental timing (Fig. 4A).

### Trus rescue experiment

To perform Trus expression rescue experiments, the UAS-Trus transgene on the 2^nd^ chromosome was combined with *Dftrus (Df(3R)BSC847)* on the 3^rd^ chromosome that was balanced with TM6B P[Dfd-GMR-nvYPF] Sb [89]. On the other hand, each of the GAL4 lines were combined with the *trus^4-15^* allele that was maintained over *TM6B P[Dfd-GMR-nvYPF] Sb* (ex. *w^1118^; ***-GAL4; trus^4-15^ / TM6B P[Dfd-GMR-nvYPF]SbTb or w^1118^;; ***-GAL4, trus^4-15^/ TM6B P[Dfd-GMR-nvYPF] Sb*). Then, *w^1118^; UAS-Trus; Dftrus/ TM6B P[Dfd-GMR-nvYPF] Sb* flies were crossed with each of the GAL4 fly line bearing the *trus ^4-15^* allele. Embryos were collected on an apple juice plate, 1st instar larvae without the nvYFP marker (*\*\*\*-GAL4>UAS-Trus* in the *trus^4-15^/Dftrus* background) were collected, transferred to regular cornmeal fly food in a 60mm dish and monitoring for developmental timing.

### RNAi

To knockdown *dilp8* or *Xrp1* in a *trus* mutant, *UAS-Dilp8RNAi* or *UAS-Xrp1RNAi* on the 2nd chromosome was combined with *Dftrus* on *TM6B P[Dfd-GMR-nvYPF]SbTb* balancer chromosome to make *w^1118^; UAS-Dilp8RNAi; Dftrus/ TM6B P[Dfd-GMR-nvYPF]SbTb* and *w^1118^; UAS-Xrp1RNAi; Dftrus/ TM6B P[Dfd-GMR-nvYPF]SbTb*. Those flies were crossed with *w^1118^;; da-GAL4, trus^4-15^ / TM6B P[Dfd-GMR-nvYPF]SbTb* to ubiquitously induce the RNAi. 1st instar larvae negative for nvYFP were transferred to 60mm culture dishes that contain regular fly food at 24-36 hours AEL for following developmental timing.

To knockdown *trus*, each GAL4 driver line was crossed with a fly line bearing the *UAS-TrusRNAi* and UAS-Dicer2 transgenes. 250-400 1st instar larvae of *nub-GAL4>UAS-TrusRNAi*, *pdm2-GAL4>UAS-TrusRNAi*, or *en-GAL4>UAS-TrusRNAi* were collected along with controls (*w^1118^* and *UAS-TrusRNAi* without a driver) and monitored for developmental timing.

### Trus localization in S2 cells

To express N-terminal EGFP-tagged Trus protein in S2 cells, the 6xHis-EGFP-HRV3C and Trus coding sequences (BDGP: RE69372) were assembled together on the *pMT-puro* vector using NEBuilder HiFi DNA Assembly (New England Biolabs). EGFP-tagged Trus expression induced with da-GAL4 *in vivo* completely rescued the *trus* mutant (*trus^4-15^/Dftrus*) phenotypes similarly to un-tagged Trus (data not shown), indicating that N-terminal EGFP tagging does not disrupt Trus function. S2 cells were transfected with the *pMT-HisEGFP-Trus-puro* plasmid and a stably transfected cell line was established by a few weeks of puromycin selection. The stably transfected S2 cells were cultured in 12-well plates and induced EGFP-Trus expression was induced with 0.7mM CuSO4. At day3, the cells were suspended in flesh media and a Concanavalin A-coated cover glass was placed in each well and left for 30 min. The wells were treated with or without 10nM Leptomycin B (InvivoGen, San Diego CA). The cover glasses were removed from the well at either 15 mins. or 115 mins., and S2 cells on each cover glass were immediately fixed with 3.2% paraformaldehyde/PBS, washed, stained with DAPI, and rinsed. Each cover glass was mounted on a slide glass facing up-side-down with DABCO/glycerol/PBS anti-fade mounting media. Edges of the cover glass were sealed with nail polish. EGFP and DAPI signals were scanned with the LSM710 laser confocal scanning microscope (Zeiss).

### Purification of protein complexes containing Trus

The coding sequence of trus from cDNA clone RE69372 was cloned into pMK33-NTAP [73], and the resulting constructs were stably transfected into *Drosophila* Kc cells using CellFectin (Thermo Fischer Scientific). The TAP-Trus fusion protein was assayed on Western blots using HRP-conjugated anti-Protein A antibody (Rockland, Gilbertsville PA) and by immunofluorescence of fixed cells using goat anti-Protein A antibody at 1:1000 followed by TRITC-conjugated anti-goat antibody (Jackson Laboratories, Bar Harbor ME) at a dilution of 1:500. Protein complexes from one liter of TAP-Trus-expressing cells were isolated following Puig *et al.*, 2001 [90], using a lysis buffer for making *Drosophila* extracts [73]. After purification using IgG-Sepharose and Calmodulin-Sepharose beads (Thermo Fisher Scientific - Invitrogen), the final eluate was precipitated with trichloroacetic acid (TCA), resolubilized in Laemmli sample buffer (BIO-RAD) and subjected to SDS-PAGE. Bands were excised, trypsinized and analyzed by MALDI (Matrix-assisted laser desorption/ionization) mass spectrometry (Cornell Institute of Biotechnology) [91].

## Acknowledgements

We thank Melissa Harrison & Kate O’Connor-Giles & Jill Wildonger for pU6-BbsI-chiRNA (Addgene # 45946; http://n2t.net/addgene:45946; RRID:Addgene_45946) and Jeffrey Lee (University of Toronto, CA) for the *pMT-puro* vector. We thank Ruth Steward for *Zfrp8* mutant flies, Bloomington *Drosophila* Stock Center (BDSC) and Vienna Drosophila Resource Center (VDRC) for other fly stocks. We thank Aidan Peterson and Myung-Jun Kim for critical reading of the manuscript. MBO thanks Pierre Leopold and the Curie Institute for providing lab space and a Mayent-Rotchild sabbatical fellowship that supported early stages of this work. Subsequent work was supported by a grant R35 GM 118029 from NIGMS to MBO.

## Author Contributions

**Conceptualization** Saeko Takada, Bonnie J. Bolkan, and Michael B. O’Connor

**Formal Analysis** Saeko Takada

**Funding Acquisition** Michael B. O’Connor

**Investigation** Saeko Takada, Bonnie J. Bolkan and MaryJane O’Connor

**Methodology** Saeko Takada and Michael B. O’Connor

**Project Administration** Michael B. O’Connor

**Resources** Saeko Takada, Bonnie J. Bolkan, MaryJane O’Connor and Michael B. O’Connor

**Supervision** Michael Goldberg and Michael B. O’Connor

**Visualization** Saeko Takada

**Writing - Original Draft Preparation** Saeko Takada and Michael B. O’Connor

**Writing – Review & Editing** Saeko Takada, Bonnie J. Bolkan, MaryJane O’Connor, and Michael B. O’Connor

---

**Fig. S1.**
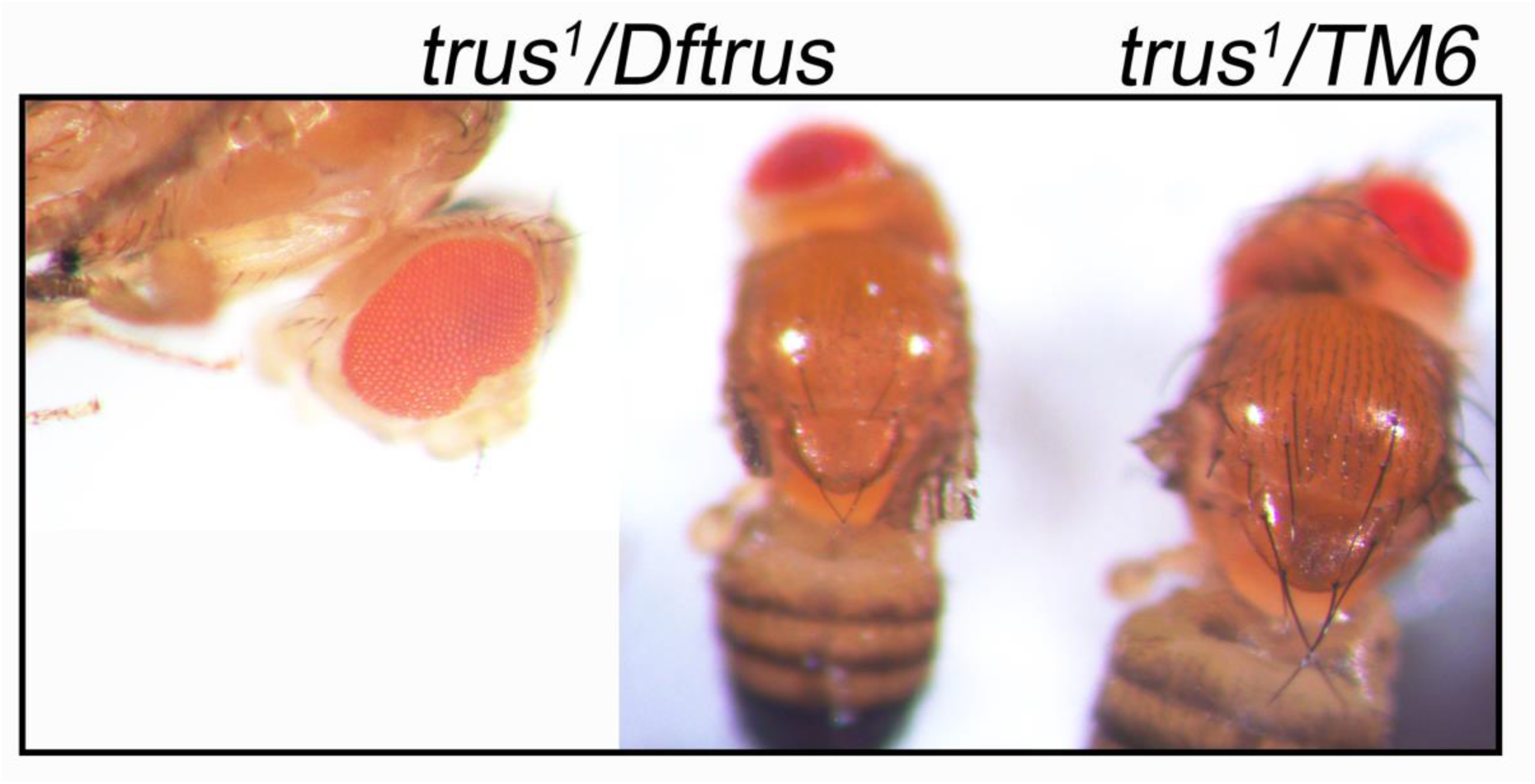
*trus^1^* mutant shows ‘Minute’ syndrome phenotype. *trus^1^* mutant (*trus^1^/Dftrus*) larvae delay development, and most of them are pre-pupal lethal. Rare escaper adults show rough/notched eyes and thin/short bristles which resemble the haplo-insufficiency ‘Minute’ syndrome that is often observed in flies carrying a mutation in one of the genes encoding ribosomal proteins.

**Fig. S2.**
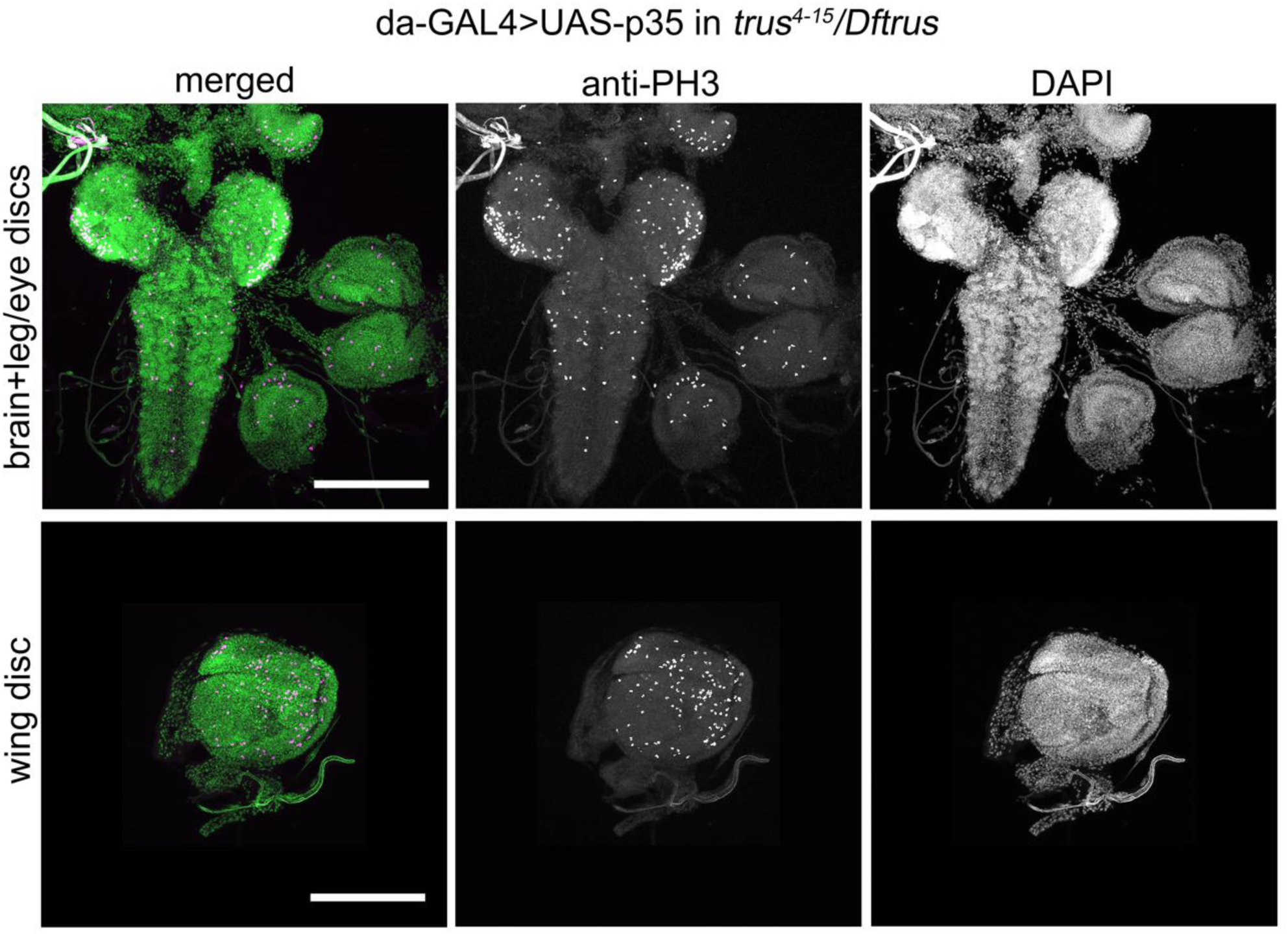
Overexpression of p35, an apoptosis inhibitor, did not rescue the defects in tissue growth and cell proliferation in *trus* mutant larvae. Representative images of brain (top row) and wing discs (bottom row) from *da>UAS-p35* in *trus^4-15^/Dftrus* third instar wandering larvae are shown. In the left column, DNA staining with DAPI (green) is merged with anti-PH3 antibody staining (magenta). Individual channels are shown in the middle and right columns. Brain, leg discs, and wing discs are smaller than those dissected from wild type (Fig. 1D) and their structures are disturbed. Scale bar, 200μm.

**Fig. S3.**
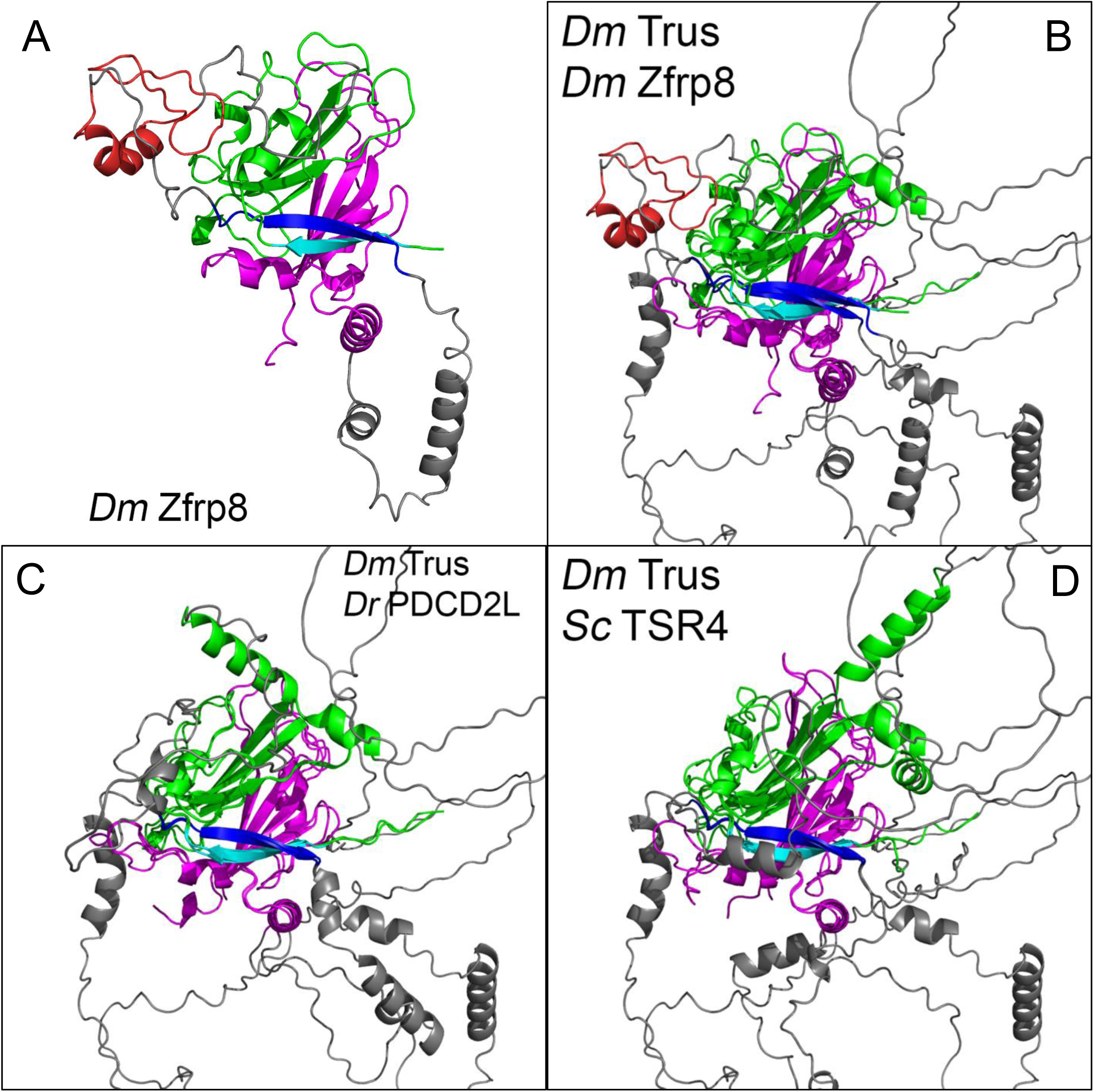
*Drosophila* Trus and its paralog Zfrp8 share a core structural module that is evolutionarily conserved. **(A)** AlphaFold structure of *Drosophila melanogaster* Zfrp8 (Accession number: Q9W1A3) with domains PDCD2_N (green and light blue), PDCD2_C (magenta), β-strand (blue) that interacts with b-strand (light blue), and the MYND-type Zinc finger (red). **(B)** Alignment of the core module of *Drosophila* Trus (*Dm*Trus) to its paralog *Drosophila* Zfrp8 (*Dm*Zfrp8). **(C)** Alignment of the core module of *Drosophila* Trus (*Dm*Trus) with Zebrafish PDCD2L (*Danio rerio* PDCD2L). **(D)** Alignment of the core module of *Drosophila* Trus (*Dm*Trus) with yeast TSR4 (*Sc*TSR4). All structures presented are predicted by AlphaFold (https://alphafold.ebi.ac.uk/), and structural alignment was performed using PyMOL (https://pymol.org/2/).

**Fig. S4.**
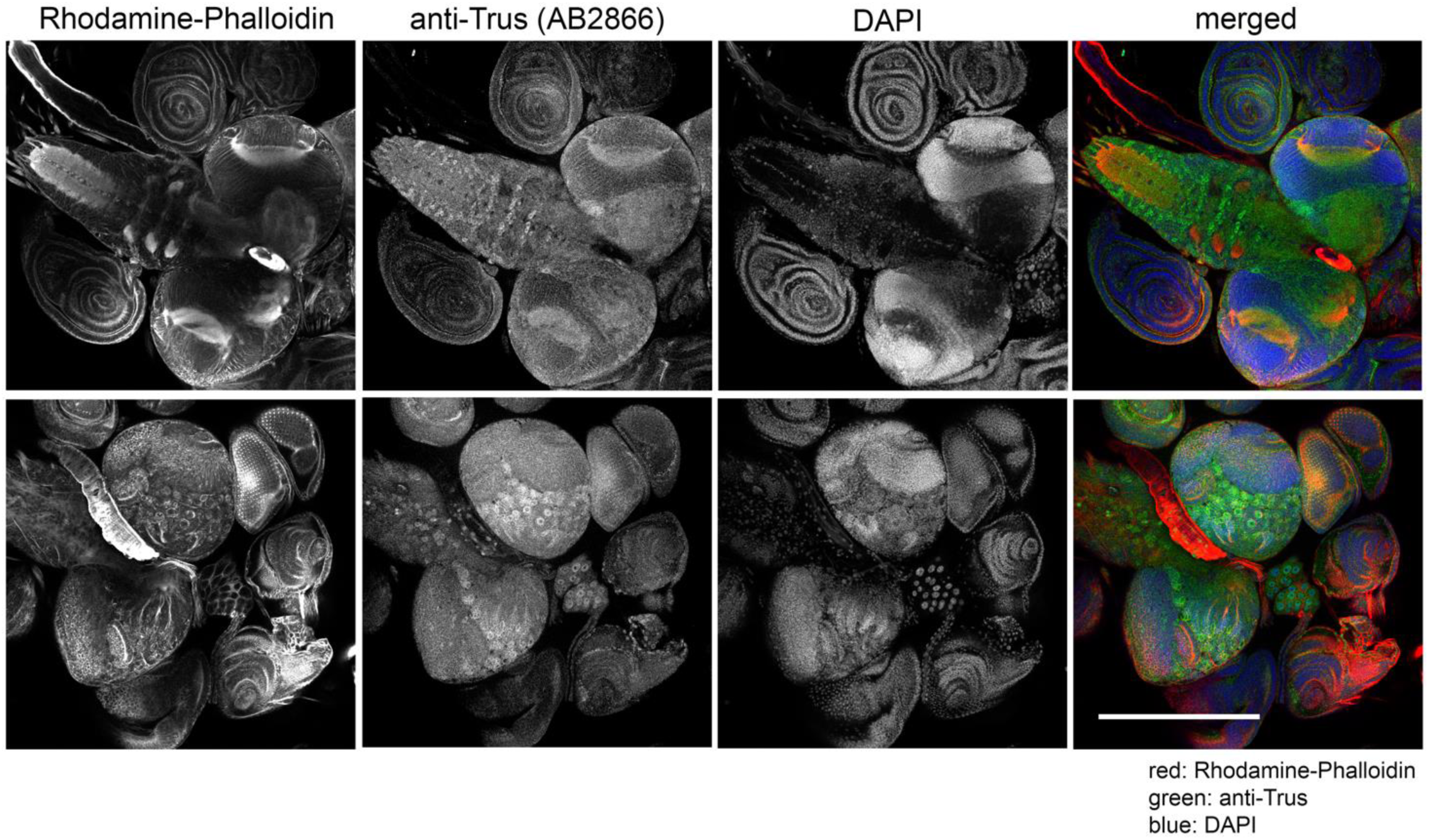
Trus protein is endogenously expressed in the central nervous system (CNS). CNS and disc tissues from *nub>TrusRNAi* 3rd instar wandering larvae were stained with affinity-purified anti-Trus antibody (green), Rhodamine-Phalloidin for F-actin (red), and DAPI for DNA (blue). Individual channels on the left and merged image on the right. Trus protein is expressed in brain lobes, the VNC, leg discs, eye-antenna discs, and the prothoracic gland.

**Fig. S5.**
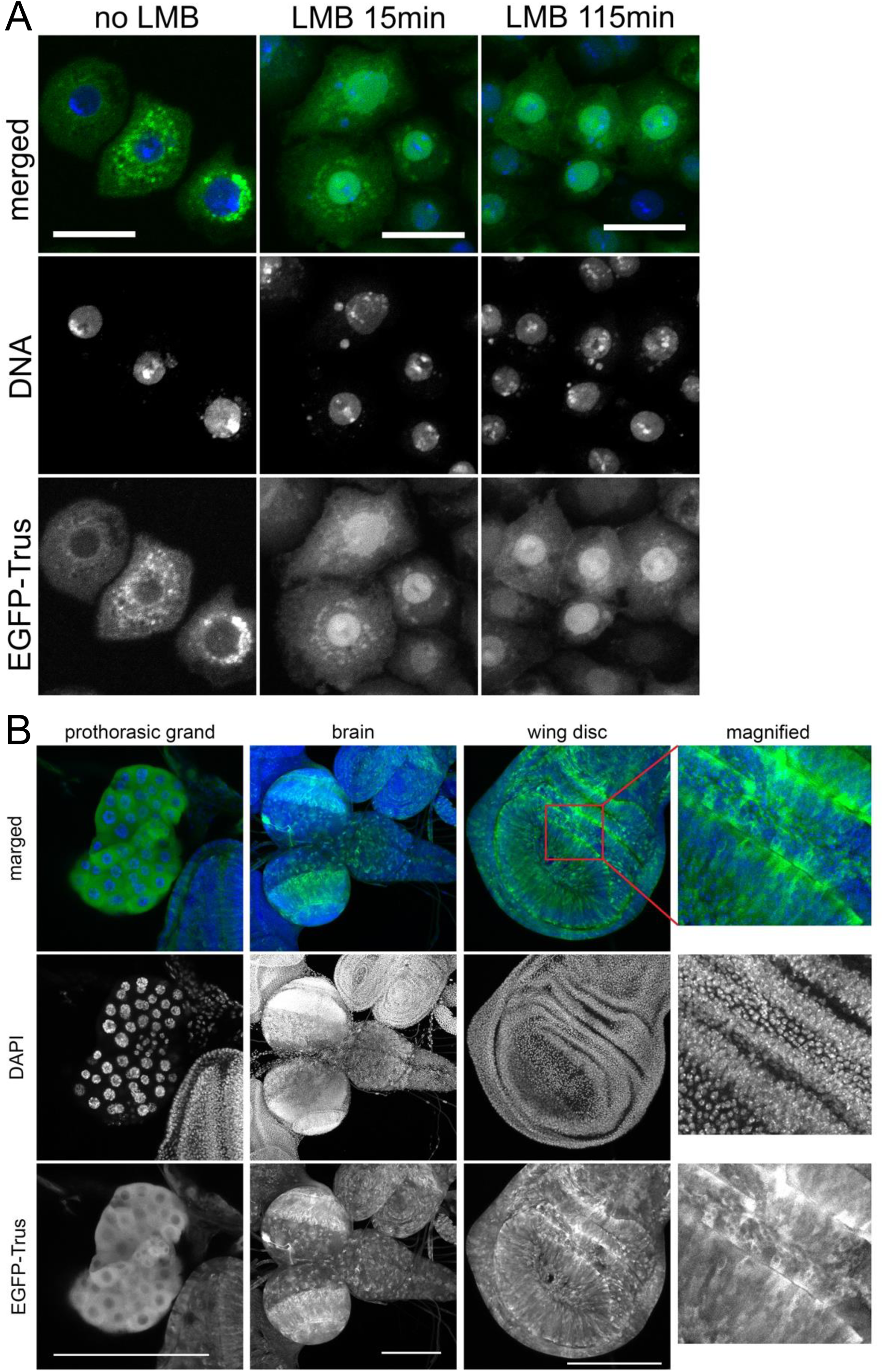
Trus localizes to the cytoplasm in cultured cells and *in vivo*. **(A)** *Drosophila* Trus localizes to the cytoplasm in S2 cells and shuttles between the nucleus and the cytoplasm in a CRM1-dependent manner. With no LB treatment (left panels), EGFP-Trus expressed in S2 cells localizes to the cytoplasm, and after treatment of the cells with Leptomycin B (middle and right panels), an inhibitor of CRM1, EGFP-Trus accumulates in the nucleus and is depleted from the cytoplasm (LMB 15min, LMB 115min). Bar: 20mm. **(B)** *da>UAS-EGFP-Trus* is ubiquitously expressed in most tissues due to *daughterless* expression. EGFP-Trus (green) staining is shown to localize in the cytoplasm of the prothoracic grand (left panels), brain lobe and ventral nerve cord (second from left panels), and the wing disc (2 panels on right). Scale bar: 200mm. Magnified images of a wing pouch region are shown in the right column.

**Fig. S6.**
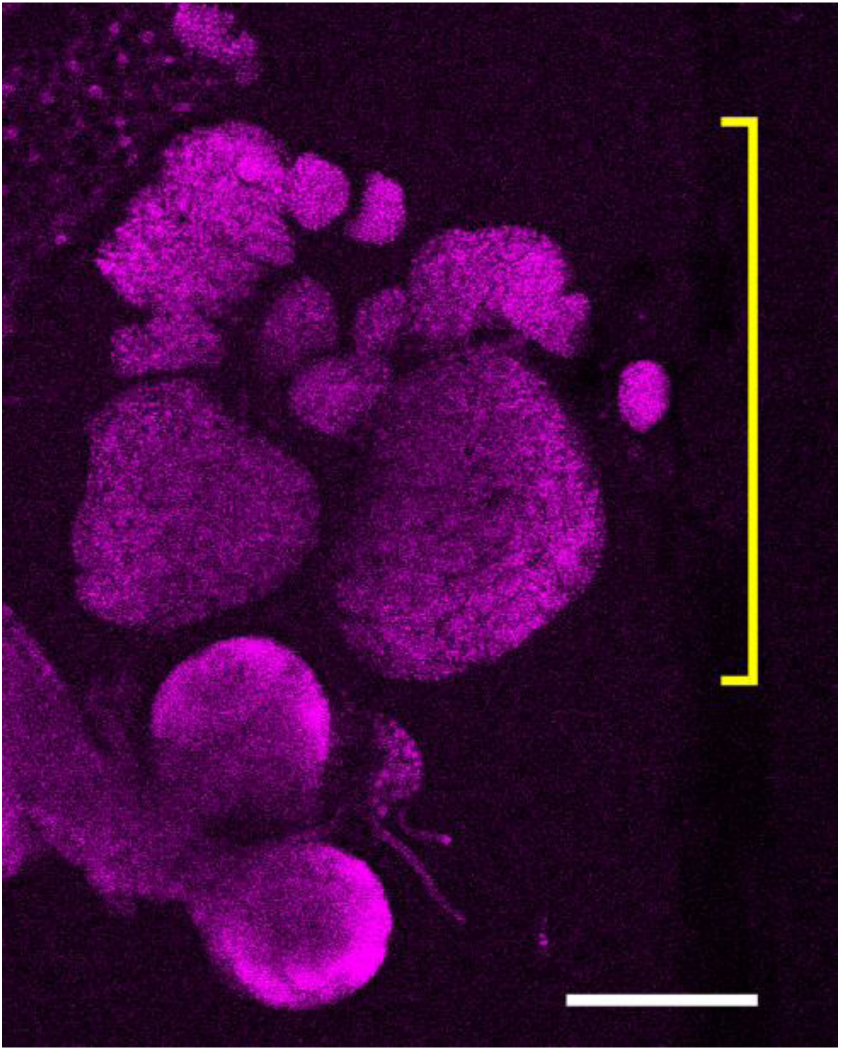
Enlarged lymph gland in trus1/trus1 larva. Lymph gland is marked with yellow bracket. Scale bar: 200µm.

**Supplementary Table 1.**
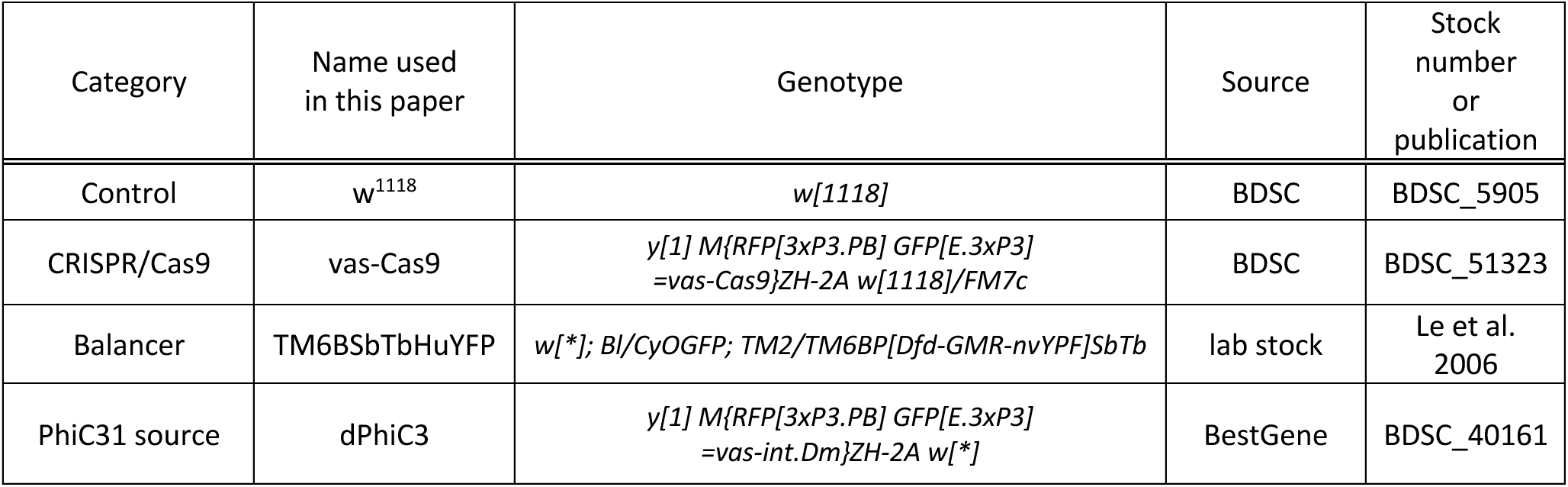
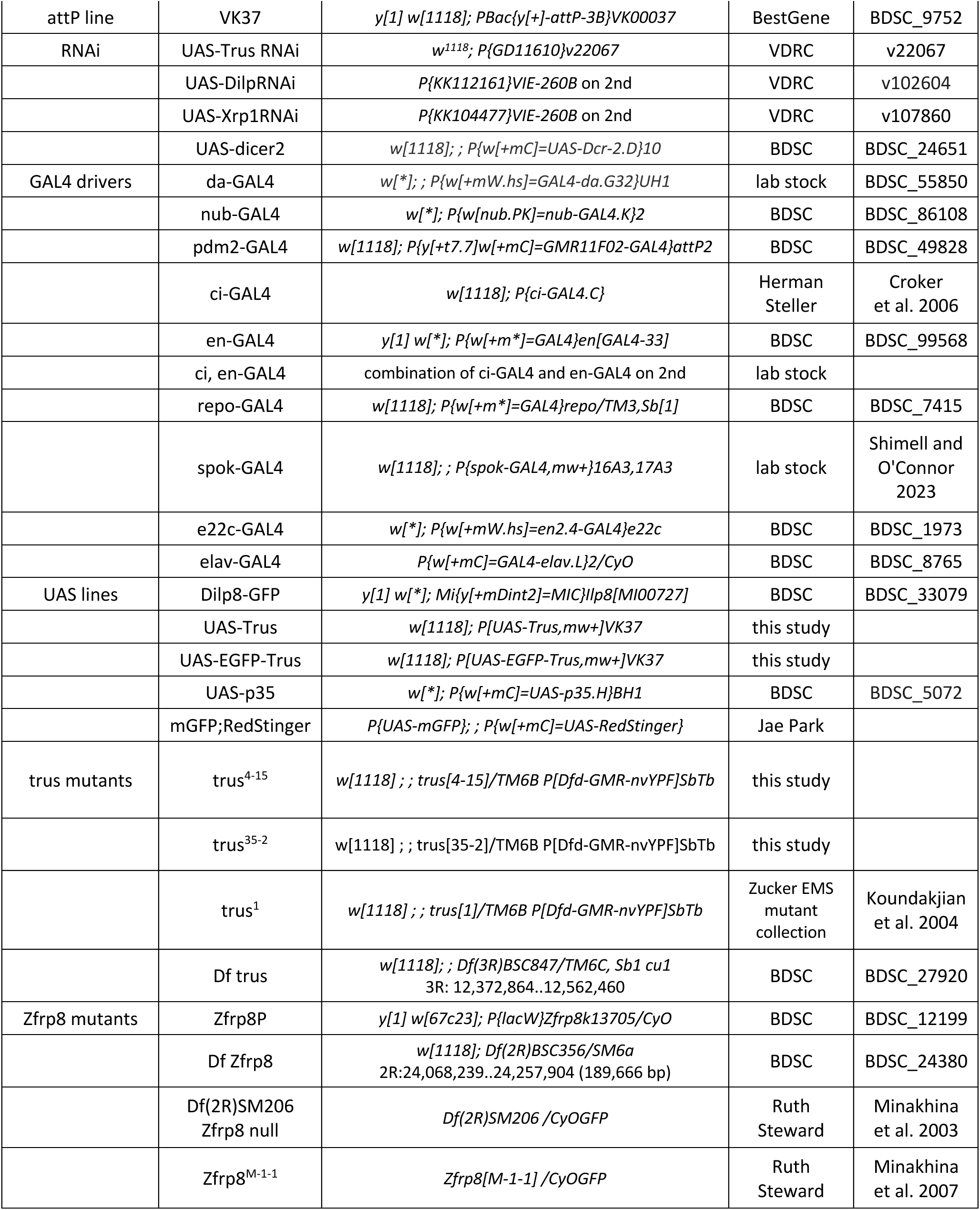
Drosophila lines that are used in this study.

**Supplementary Table 2.**
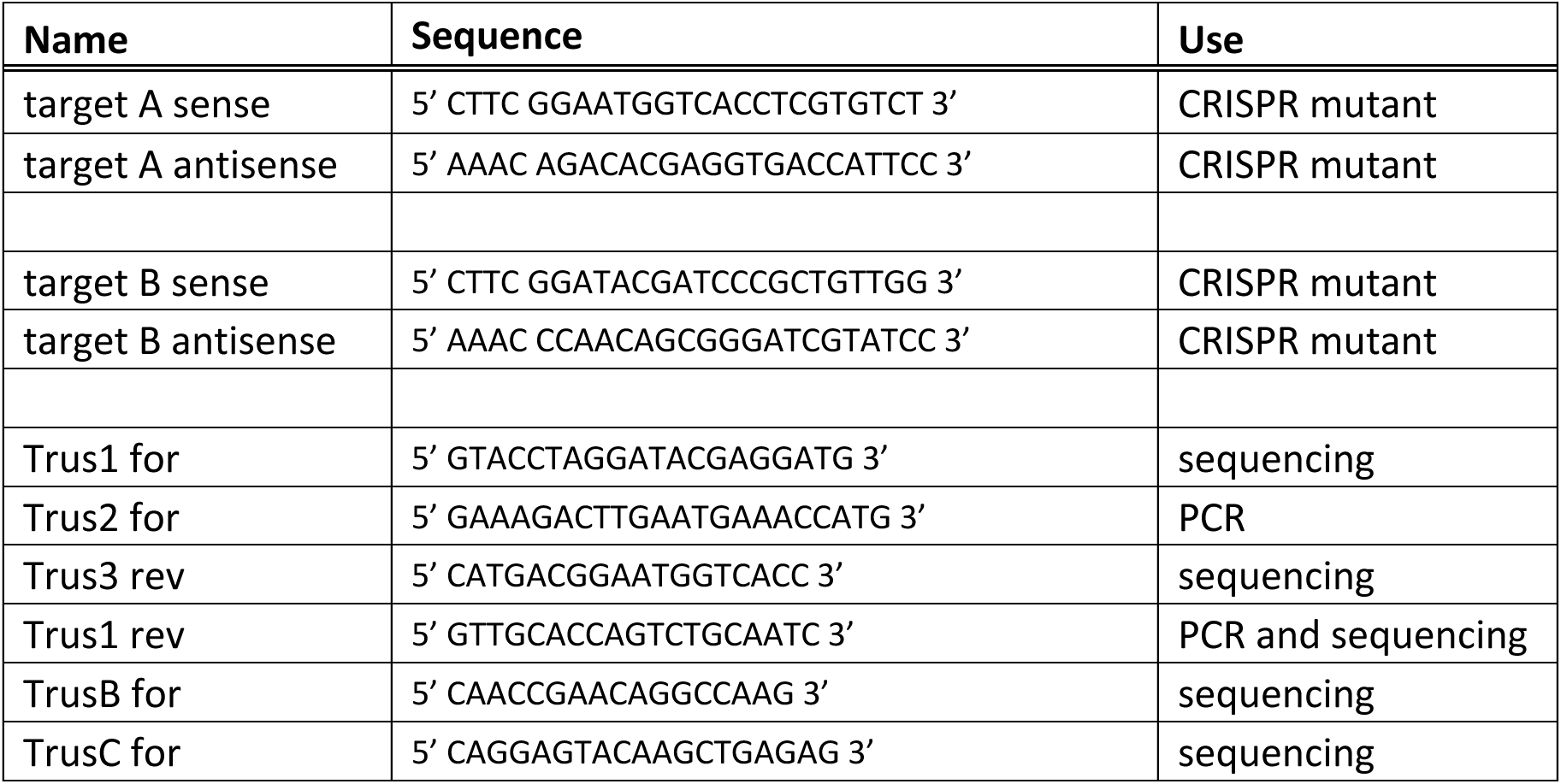
DNA oligos that are used for production of CRISPR/Cas9 trus mutants.

